# Cisplatin-triggered sensory dysfunction in NER-defective *C. elegans* involves GABAergic and cholinergic signaling

**DOI:** 10.64898/2026.11.06.764850

**Authors:** Johannes Krautstrunk, Emre Inan, Andre Potthoff, Ann-Kathrin Weishaupt, Thorsten Wachtmeister, Julia Bornhorst, Simone Prömel, Gerhard Fritz

**Affiliations:** Heinrich-Heine-University Düsseldorf, Medical Faculty, Institute of Toxicology, Moorenstraße 5, 40225 Düsseldorf, Germany; University of Wuppertal, Faculty of Mathematics and Natural Sciences, Food Chemistry with Focus on Toxicology, Gaußstraße 20, 42119 Wuppertal, Germany; Heinrich Heine University Düsseldorf, Genomics & Transcriptomics Laboratory, Biomedical Research Centre (BMFZ), Universitätstraße 1, 40225 Düsseldorf, Germany; Heinrich-Heine-University Düsseldorf, Department of Biology, Institute of Cell Biology, Universitätstraße 1, 40225 Düsseldorf, Germany

**Keywords:** *Caenorhabditis elegans*, Cisplatin, DNA damage, DNA repair (NER), sensory neurotoxicity, GABAergic neurotransmission

## Abstract

Cisplatin (CisPt) is a widely used anticancer drug that causes neurotoxicity as a dose-limiting adverse effect. DNA damage contributes to CisPt-induced neurotoxicity, yet the molecular mechanisms involved remain unclear. Here, we aim to identify pathophysiological mechanisms contributing to DNA damage-driven development of sensory neuronal dysfunction using the 3R-compliant model organism *Caenorhabditis elegans*. To this end, we employed the nucleotide excision repair (NER)-deficient *ercc-1* mutant, which shows enhanced CisPt-induced sensory dysfunction as monitored by the chemotaxis assay, to elucidate drug-induced alterations in neurotransmission and transcriptome.

We found that CisPt reduces gamma-aminobutyric acid (GABA) levels in the nematode and that GABA supplementation rescues *ercc-1* mutants from CisPt-evoked sensory neurotoxicity, pointing to a neuroprotective function of GABA in the context of our study. Moreover, CisPt impairs the morphological integrity of GABAergic neurons particularly in *ercc-1* mutants, which again is rescued by GABA substitution. Conversely, the cholinergic agonist carbachol impairs sensory function and exacerbates the neurotoxic effects of CisPt in *ercc-1* mutants, indicating detrimental effects resulting from cholinergic excitation. Transcriptome analysis revealed that CisPt treatment stimulates a broard transcriptional response, including pathways involved in the regulation of neuronal communication, including acetylcholine-mediated neurotransmission, with pronounced effects occurring in the absence of ERCC-1. Based on these finding we suggest impaired GABAergic signaling and the loss of neuronal integrity GABAergic neurons as well as excessive cholinergic excitation as key mechanisms linking CisPt-induced DNA damage in NER defective nematodes to sensory dysfunction. Hence, targeting the balance between GABAergic inhibition and cholinergic excitation is suggested to counteract sensory neurotoxicity triggered by CisPt.

**Highlights:**

- Deficiency of the NER factor ERCC-1 aggravates sensory dysfunction following CisPt treatment.
- CisPt-induced DNA damage-driven sensory neurotoxicity coincides with neuronal beading.
- GABA substitution restores sensory function and morphological alterations of *ercc-1* mutants.
- Carbachol further increases CisPt-mediated neurotoxicity in *ERCC-1* defective nematodes.
- CisPt treatment causes substantial changes in the transcriptome, preferably in *ercc-1* mutants.

## 1. Introduction

Cisplatin (CisPt) is part of the first line treatment regimen for the therapy of numerous malignant diseases, including lung, bladder and ovarian carcinoma (1). Its anticancer efficacy rests on the formation of DNA adducts, especially DNA intrastrand-crosslinks (i.e. 1,2-d(GpG) and 1,2-d(ApG)) as the most frequent type of DNA adduct (80-90%) (2). In addition, DNA interstrand-crosslinks represent another important type of DNA damage formed by CisPt (3-5%). CisPt-induced DNA crosslinks cause a distortion of the DNA helix, thereby hampering DNA replication and transcription. Nucleotide excision repair (NER), which subdivides into global genome nucleotide excision repair (GG-NER) and transcription-coupled repair (TC-NER) (3, 4), is of utmost relevance for the removal of CisPt-formed DNA adducts. After damage recognition, the excision of the DNA adducts is mediated by the ERCC1/XPF complex (5’ endonuclease) and XPG (3’ endonuclease), followed by gap filling (i.e., repair synthesis) and ligation (5). Correspondingly, lack of NER-associated DNA repair promotes DNA damage-related degenerative processes as well as aging (5-9).

In clinical practice, normal tissue damage caused by CisPt limits the maximal applicable dose. Apart from acute nephrotoxicity (10) and cardiotoxicity (11), chemotherapy-induced neuropathy (12) is a frequent and long-lasting adverse effect of CisPt-based anticancer therapy occurring in up to 90 % of patients (13). Chemotherapy-induced peripheral neuropathy (CIPN) is a major type of neurotoxicity caused by platinating agents that predominantly affects sensory neurons, thereby causing allodynia, numbness, paresthesia, spontaneous pain and hypersensitivity to cold stimuli (14, 15). These symptoms can persist for a long period of time even after cessation of the anticancer treatment and substantially reduce the quality of life of cancer patients (12). Sensory disturbances have been linked to the accumulation of CisPt in the dorsal root ganglions (DRGs) (16), which harbor the cell bodies of primary sensory neurons and represent a crucial site for the transmission of peripheral sensory input through the dorsal horn of the spinal cord to the central nervous system (17). Healthy sensory perception relies on proper signaling, including inhibitory signaling of gamma-aminobutyric acid (GABAergic) neurons for regulating nociceptive input and, moreover, preventing excessive signal transmission to the brain (18-20). Disruption of these inhibitory circuits contributes to neuropathic pain. In case of mechanical allodynia, impairment of inhibitory signaling in the dorsal horn results in pain hypersensitivity (21, 22). Besides sensory toxicity, CisPt-induced neurotoxicity may also involve motor disorders and cognitive impairment (23, 24). The molecular mechanisms contributing to CisPt-induced neurotoxicity are still unclear and effective treatment or prevention is not availabe. Having in mind that DNA damage accelerates aging (25) and various diseases, including neurodegeneration (26-29), it is feasible that DNA damage evoked by CisPt-based anticancer therapy also contributes to the onset of CIPN (30, 31).

*Caenorhabditis elegans* (*C. elegans*) is a well-established 3R-conform model in neurobiology, toxicology and pharmacology (32-34). It is useful to study mechanims of DNA damage response (35) and DNA damage-driven diseases (35, 36), including CisPt-induced neurotoxicity (37-39). Many functions that ensure genetic stability are well-conserved in *C. elegans*, including mechanisms of DNA repair (35, 38, 40), making the nematode a meaningful model organism to investigate the impact of genetic instability on aging (6, 9). For instance, it has been demonstrated that NER protects the nematode from UV-induced DNA damage (41, 42) and neurodegeneration (36). Of note, *C. elegans* is also useful to study adverse effects of platinating anticancer agents (34, 39). Moreover, CisPt-induced DNA crosslinks exacerbate CisPt-induced neuropathy, including sensory dysfunction, in NER deficient *C. elegans* strains (38, 43).

Here, we aim to further investigate the role of unrepaired DNA-damage in CisPt-induced neurotoxicity by comparatively analyzing the CisPt response of wildtype and NER-deficient *ercc-1* mutants. Our previous work demonstrated that ERCC-1 defective nematodes are more susceptible to CisPt-induced sensory dysfunction as reflected by a limited chemotaxis towards the chemoattractant diacetyl. This effect likely results from functional neuronal impairment rather than apoptosis of chemosensory neurons (43). Since a healthy sensory function critically depends on accurate neurotransmission, we hypothesize that unrepaired CisPt-induced DNA damage may impair neuronal signalling by disturbing neurotransmitter homeostasis. To scrutinize this, neurotransmitter levels were comparatively analyzed in wildtype and *ercc-1* muant animals to identify potential imbalances. In addition, we performed transcriptomic profiling by RNA sequencing (RNA-seq) to characterize alterations in molecular pathways associated with CisPt-induced neuronal insult in wildtype nematodes and ERCC1-defective mutants. Thereby, we aim to elucidate molecular mechanisms involved in CisPt-induced sensory neurotoxicity.

## 2. Materials and Methods

### 2.1. *C. elegans* strains and nematode maintenance

For this study, we used the following *C. elegans* strains: N2 [wildtype], CB156 [*unc-25(e156) III*], CB382 [*unc-49(e382) III*], CHS1016 [*gbb-2(yum1169) IV; gbb-1(yum1168) X*], CX3260 [*odr-10::GFP + lin-15(+)*], EG1285 [*unc-47p::GFP + lin-15(+)*], GG201 [*ace-2(g72) I; ace-1(p1000) X*], PR1162 [*cha-1(p503) IV*], PS8727 [*lgc-37(sy1492) III*], PS8743 [*lgc-36(sy1502) V*], RB1124 [*exp-1(ok1131) II*], RB1559 [*acr-2(ok1887) X*], RM2576 [*cho-1(tm373) IV*], TG1663 [*ercc-1(tm1981) I*], TJ1 [*cep-1 (gk138) I*], VC381 [*atm-1(gk186) I*], VC670 [*gar-3(gk337) V*] and VC728 [*atl-1(ok1063) V*]. The strains were provided by the Caenorhabditis Genetics Center, funded by the NIH Office of Research Infrastructure Programs (P40 OD010440), or the *C. elegans* Gene Knockout Project at the Oklahoma Medical Research Foundation, which was part of the International *C. elegans* Gene Knockout Consortium (44). The strains CX3260, EG1285 and VC381 were crossed with TG1663 to introduce an *ercc-1* defective genetic background, leading to the generation of the following novel strains: JK01 [*odr-10::GFP + lin-15(+); ercc-1(tm1981) I*], JK03 [*unc-47p::GFP + lin-15(+); ercc-1(tm1981) I*] and JK04 [*atm-1(gk186) I; ercc-1(tm1981) I*]. The genotypes were confirmed by genomic polymerase chain reaction. All strains were maintained at 20°C on NGM plates seeded with *Escherichia coli* strain OP50 (45).

### 2.2. Synchronization of nematodes

For all experiments, *C. elegans* populations were age-synchronized using a standard alkaline hypochloride bleaching protocol (45). Briefly, an isogenic population containing mainly well-fed gravid adult worms was rinsed off NGM plates and collected in 1 mL S-buffer (100 mM NaCl, 5.7 mM K_2_HPO_4_, 44.1 mM KH_2_PO_4_). A bleaching solution was added to an end concentration of 1% NaClO and 0.4 M NaOH and incubated until the adult worms started to dissolve leaving only the eggs intact. The bleaching solution was removed from the tubes through three washing steps with S-buffer. The isolated eggs were placed on fresh NGM plates and incubated at 20°C for 48 h until reaching late L4/ young adult stage.

### 2.3. Compound treatment

In this study, age-synchronized *C. elegans* populations were treated both in solid and liquid culture. For solid culture treatment, the worms were kept on prepared nematode growth medium (NGM) agar containing CisPt. CisPt (Sigma-Aldrich GmbH, Darmstadt, Germany) was dissolved in sterile saline (150 mM NaCl) and added to freshly autoclaved NGM to the desired concentration. Solidified treatment plates also contained 0.01% ampicillin and were seeded with OP50 (*xu363*) bacteria containing the L4440 plasmid (46). The worms were rinsed off the plates daily, washed at least three times with S-buffer and were transfered onto fresh treatment plates.

For liquid culture treatment, the worms were suspended in S-medium containing 1% bovine serum albumin and OP50-1 (OD_600_ 2.5) according to (45). CisPt was dissolved in sterile saline and was added to the liquid medium to the desired concentration. For neurotransmitter treatment, the neurotransmitter were dissolved at the desired concentration directly in S-medium. Every day, the worms were washed with S-buffer at least three times and suspended in fresh medium.

### 2.4. UV treatment

Age-synchronized worms were irradiated in solid culture on NGM agar plated containing 0.01% ampicillin, seeded with OP50 (*xu363*) bacteria containing the L4440 plasmid (46). Animals were exposed to ultraviolet C (UV-C) irradiation using the UV-C Crosslinker Biolink BLX-254. Following irradiation, the worms were rinsed off the plates daily, washed at least three times with S-buffer and were transfered onto fresh plates.

### 2.5. Chemotaxis assay

Dysfunction of sensory neurons was assessed using the 4-quadrant method of the chemotaxis assay (47). Briefly, 10 cm diameter chemotaxis plates (2 % (w/v) agar, 5 mM KPO_4_ buffer, 1 mM CaCl_2_, 1 mM MgSO_4_) were divided into four equal quadrants. Opposing quadrants were marked with the substances “A” (attractant/odorant) or “C” (control/ethanol) in a distance of 3.5 cm from the center of the plate. The center was marked with a 1 cm diameter circle to define the starting area. In the quadrant marked with “A”, 1 µl of the attractant solution (0.5% diacetyl diluted in 96% ethanol) was applied as well as 1 µl of 1 M NaN_3_ to immobilize the worms upon arrival. In the quadrant marked with “C”, 1 µl of the solvent (96% ethanol) and 1 µl 1 M NaN_3_ was applied. Treated worms were washed four times with 1 ml S-buffer and once with 1 ml sterile water. To start the assay, >100 worms per experimental condition were applied in the center of the prepared plates. After removing the excess water, the plates were incubated at 20°C for one hour. Afterwards, the number of worms present in the attractant “A” and control “C” quadrants was counted. Worms that remained in the starting area were excluded from analysis. Chemotactic behavior was quantified using the chemotaxis index (CI):

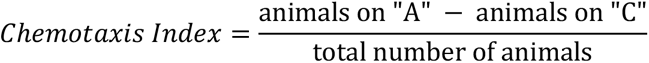

### 2.6. Swimming assay

Swimming behavior was assessed by monitoring the number of thrashes in liquid medium. Prior to the assay, worms treated in liquid medium were allowed to recover on NGM plates seeded with OP50 for >1 h to minimize potential acute effects of liquid culture and handling. For each experimental condition, ten worms were indviually transferred into a 20 µl S-buffer droplet on a microscope slide. Thrashes were counted for 30 seconds under a stereomicroscope.

### 2.7. Evaluation of neuron morphology

To evaluate neuron morphology/degeneration of *C. elegans*, age-synchronized drug-treated worms carrying green fluorescent protein (GFP) reporter transgenes specifically expressed in either chemosensory AWA neurons (*odr-10::GFP*) or GABAergic neurons (*unc-47p::GFP*) were analyzed by fluorescence microscopy. Pseudo Z-stack images of GFP-labeled neurons were taken at a 400x magnification with an Olympus CKX41 microscope in the GFP channel using an Olympus XM10 camera. Neuronal degradation was quantified by counting the number of bead-like outgrowths along the visible course of neurons (48). To take into account differences in the length of the individual neurons analyzed, chemosensory AWA neurons were scored by the number of outgrowths per neuron while longer GABAergic body neurons were scored by counting ̴ 200-300 µm of anterior D-type ventral neuron and caltulating the outgrowths per µm. Neurons were classified as undergoing cell death only when the neuron was fragmented / with no visible connection between adjacent beads.

### 2.8. RNA-seq and bioinformatic analyses

RNA was isolated from treated worms (100 µg/ml CisPt, 24 h, solid culture treatement) using the RNeasy-/ Shredder Kit (Qiagen) and quality-assessed by capillary electrophoresis (Fragment Analyzer, DNF-471 Total RNA High Sensitivity Assay; Agilent Technologies, CA, USA). 200 ng total RNA were used to prepare libraries using the QuantSeq 3′ mRNA-seq V2 Kit with UDI/UMI Second Strand Synthesis Module (Lexogen, Vienna, Austria) following the manufacturer’s protocol. Following bead-purification and library normalization, sequencing was performed at the Center for Biological-Medical Research (BMFZ) (Heinrich Heine-University Düsseldorf, Germany) on the NextSeq2000 platform (Illumina, CA, USA) in single-end mode (101 bp). BCLConvert (v4.2.7) was used for BCL-to-FASTQ conversion, adapter trimming, and demultiplexing. Subsequent bioinformatic analyses were conducted using the CLC Genomics Workbench (v24.0.1; QIAGEN, Venlo, Netherlands). Unique molecular identifier (UMI) sequences were extracted and incorporated to read headers, reads lacking valid UMI information were excluded from further analysis. Quality control included adapter trimming and low quality base (Q < 13; maximum two ambiguous nucleotides) removal. Filtered reads were mapped to the *Caenorhabditis elegans* reference genome (WBcel235). Samples were grouped into four biological replicates per experimental condition and compared using the CLC RNA-seq analysis workflow. Raw count data were analyzed with *DESeq2* v.1.38.3 (49) using a multifactorial model incorporating genotype, treatment and their interaction. To account for large variance in fold changes of low-abundance genes, we applied *ashr* shrinkage estimation (50). Subsequently, sample distribution and variance sources were assessed through principal component analysis. Statistical significance of resulting differentially expressed genes (DEGs) was determined using false discovery rate (FDR)-adjusted p-values. DEGs were considered as biologically relevant at significance level p(adj) ≤ 0.01 and at fold change ≥ 2.

Functional enrichment analysis was performed using the Database for Annotation, Visualization and Integrated Discovery (DAVID) (version 6.8; DAVID Bioinformatics Resources (51, 52); assessed on 08.04.2026). The functional annotation clustering tool was applied to identify overrepresented biological themes among biological relevant DEGs. Gene Ontology (GO) terms, Kyoto Encyclopedia of Genes and Genomes (KEGG) pathways and Reactome pathways were included as annotation categories. Statistical significance was assessed using the modified Fisher’s exact test (EASE score) and annotation clusters with an EASE threshold of p ≤ 0.05 were considered as significant.

### 2.9. Aldicarb paralysis assay

Sensitivity to paralysis induced by the carbamate-type insecticide aldicarb was monitored to analyze cholinergic function (53). After treatment with CisPt, age-synchronized worms were allowed to recover on NGM plates seeded with OP50 for >1 h to minimize potential acute effects of liquid culture and handling. Afterwards, 21 animals per experimental condition were transferred onto 3.5 cm diameter NGM plates containing 15 µM aldicarb and seeded with OP50, prepared prior to the experiment. Animals were scored for paralysis at time intervals of 30 min. The worms were considered paralyzed if they failed to exhibit spontaneous movement in response to gentle stimulation with an eyelash pick three times.

### 2.10. Neurotransmitter quantification

Neurotransmitter levels of serotonin, dopamine, GABA and acetylcholine were determined using a quantitative liquid chromatography-tandem mass spectrometry-based method. Samples were prepared by treating worms with CisPt on solid culture for 72 h. ∼1500 worms per condition were rinsed off the plates, washed three times using 85 mM NaCl, frozen in liquid nitrogen and stored at -80°C until analysis. Liquid chromatography-tandem mass spectrometry was performed as described before (54).

### 2.11. Statistical analysis

Statistical analysis was performed using GraphPad Prism 6. For statistical evaluation, one-way ANOVA with post-hoc tests for multiple comparision or students t-test were used for statistical evaluation. P values ≤ 0.05 (p ≤ 0.05) were considered as statistically significant differences between experimental groups.

## 3. Results

### 3.1. CisPt-induced sensory toxicity in DNA repair- and DDR-mutants

The chemotaxis assay was employed to comparatively investigate the impact of CisPt on sensory functions of wildtype and *ercc-1* mutants of *C. elegans*. Dose response analyses revealed a substantially higher sensory dysfunction in CisPt treated *ercc-1* mutants as compared to wildtype nematodes, with a low CisPt concentration of 10 µg/ml reducing the chemotaxis index (CI) already by about 50 % in the NER-defective mutant **(Fig. 1A)**. Noteworthy, swimming activity of *ercc-1* mutants was only minorly affected by such low CisPt concentration **(Fig. 1B)**, suggesting a deleterious effects of CisPt preferentially on sensory neurons of the nematode rather than motor neurons. Noteworthy, the lack of functional DNA damage response (DDR)-related factors *cep-1* (human p53 ortholog), *atm-1* (human ATM ortholog) and *atl-1* (human ATR ortholog) did not impact CisPt-induced sensory functions as compared to wildtype control animals **(Fig. 1C)**. Yet, *atm-1* mutants revealed a moderate reduction in their basal CI in the absence of CisPt **(Fig. 1C)**, indicating that ATM-1-regulated signaling influences sensory functions of the nematode under basal situation. To investigate the impact of ATM-1 on CisPt-induced sensory functions against a genetic *ercc-1* mutant background, a *ercc-1;atm-1* double mutant was generated. The *ercc-1;atm-1* double mutant exhibited a CisPt response that was comparable to that of the *ercc-1* single mutant **(Fig. 1D)**, indicating no further additional chemotaxis defect by lack of ATM-1. This finding identifies lack of ERCC-1 dependent DNA repair of CisPt-induced DNA damage, rather than canonical ATM-1-regulated DDR-related signaling as a major determinant of the high susceptibility of *ercc-1* mutants to CisPt-induced sensory impairment.

**Fig. 1:**
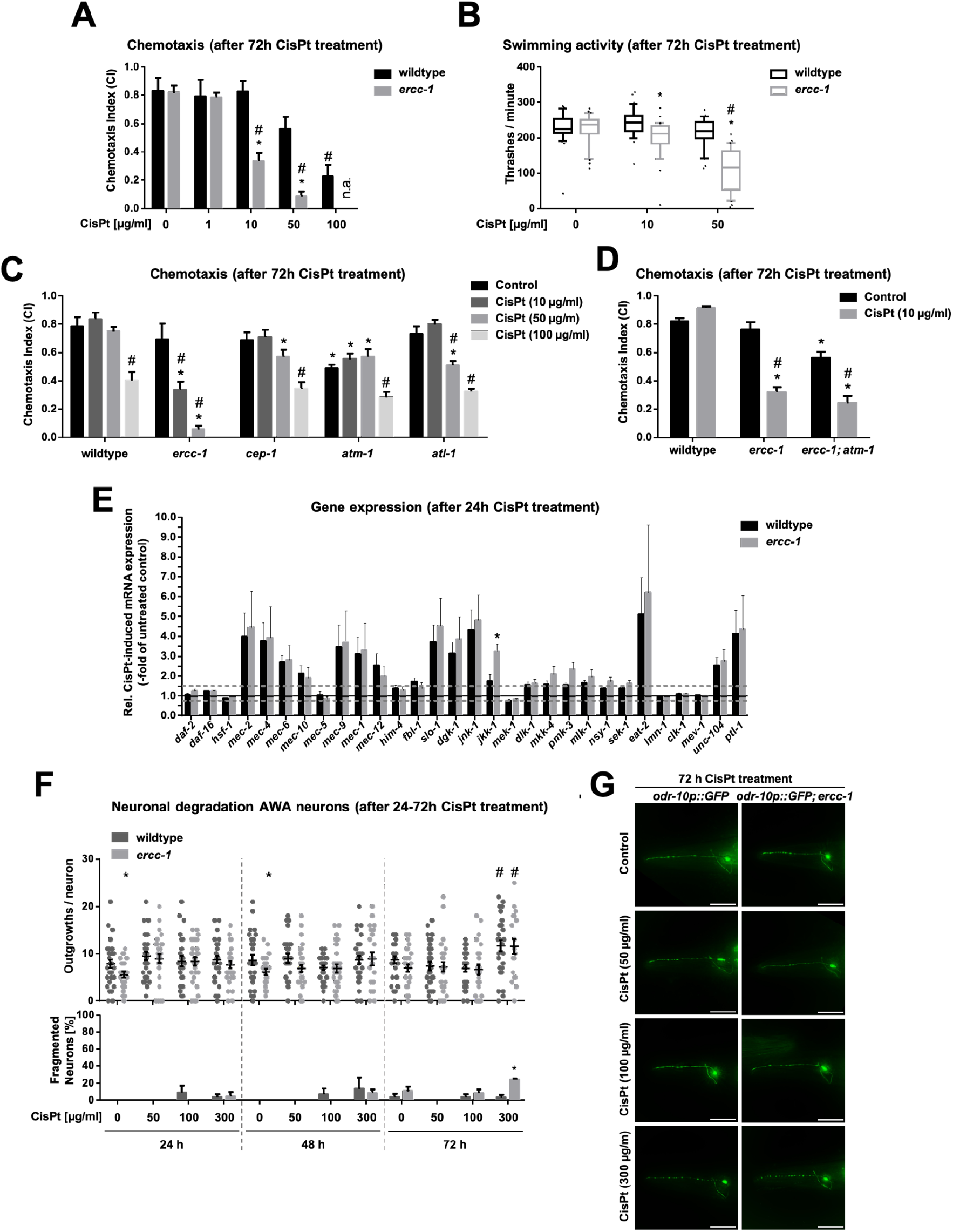
Low concentration of CisPt strongly impairs chemotaxis of *ercc-1* mutants. **A:** Chemotaxis analysis of wildtype and *ercc-1* mutated animals after 72 h of CisPt treatment (0-100 µg/ml). Data shown are the mean + SEM of three independent experiments (N ≥ 100 nematodes per experimental condition and experiment). *, p ≤ 0.05 (wildtype vs. mutant; students t-test); ^#^, p ≤ 0.05 (control vs. CisPt-treatment; One-Way ANOVA). **B:** Swimming activity of wildtype and *ercc-1* mutated animals was analyzed after 72 h of CisPt treatment (0-50 µg/ml). Data shown as boxplots with median (center line) box (25^th^-75^th^ percentiles) and whiskers (10^th^-90^th^ percentiles) of three independent experiments (N = 10 nematodes per experimental condition and experiment). *, p ≤ 0.05 (wildtype vs. mutant, students t-test); ^#^, p ≤ 0.05 (control vs. CisPt-treatment, One-Way ANOVA). **C:** Chemotaxis analysis of wildtype and DDR mutant animals (*cep-1, atm-1, atl-1*) after 72 h of CisPt treatment (0-100 µg/ml). Data shown are the mean + SEM of three independent experiments (N ≥ 100 nematodes per experimental condition and experiment). *, p≤ 0.05 (wildtype vs. mutant, students t-test); ^#^, p ≤ 0.05 (control vs. CisPt-treatment, One-Way ANOVA). **D:** Chemotaxis analysis of wildtype and *ercc-1;atm-1* double mutants after 72 h of CisPt treatment (0-100 µg/ml). Data shown are the mean + SEM of three independent experiments (N ≥ 100 nematodes per experimental condition and experiment). *, p ≤ 0.05 (wildtype vs. mutant, students t-test); ^#^, p ≤ 0.05 (control vs. CisPt-treatment, One-Way ANOVA). **E:** Relative CisPt-induced mRNA expression data obtained from RNA-seq analysis of wildtype nematodes and *ercc-1* mutants after 24 h CisPt treatment (100 µg/ml) Data shown are mean + SEM from N = 4 biological replicates per experimental condition. *, p ≤ 0.05 (wildtype vs. mutant, students t-test). **F,G:** Quantification of neuronal damage (beading and cell death) of chemosensory AWA neurons in wildtype and *ercc-1* mutated nematodes after 24 h-72 h of CisPt treatment (0-300 µg/ml) **(F)**, with representative images **(G)**. Scale bar = 50µm. Quantitative data shown are from n = 3 independent experiments, with at least 8 images being analyzed per experimental condition.*, p ≤ 0.05 (wildtype vs. mutant, students t-test); ^#^, p ≤ 0.05 (control vs. CisPt-treatment, One-Way ANOVA).

Data obtained from our previous work showed that CisPt-induced sensory dysfunction is not accompanied by neuronal apoptotic or necrotic cell death (43). Correspondingly, we assume that functional impairment rather than neuronal loss causes sensory dysfunction. To further investigate possible mechanisms of CisPt-mediated sensory dysfunction in *ercc-1* mutants, we assessed molecular and morphological alterations associated with neuronal aging. Using a selection of genes previously linked to neuronal aging (48), we observed an increased expression of some of these genes in CisPt-treated wildtype and *ercc-1* mutant nematodes. Following CisPt-exposure, both *ercc-1* and wildtype animals revealed an upregulation in the mRNA expression of the mechanosensory genes *mec-1, mec-2, mec-4, mec-6, mec-9* and *mec-12*, as well as of genes involved in the regulation of neuronal signaling and stress response, including *slo-1, dgk-1, jnk-1, jkk-1, eat-2, unc-104* and *ptl-1* **(Fig. 1E).** Notably, *ercc-1* mutants displayed a significantly increased expression of c-Jun N-terminal kinase *jkk-1 as* compared to wildtype animals **(Fig. 1E)**, indicating a stronger activation of JKK-1-regulated stress signaling by CisPt in the NER mutant, potentially contributing to its increased neuronal vulnerability.

Morphological alterations of sensory AWA neurons were investigated by counting the number of outgrowths as surrogate marker of age-dependent neurodegeneration in NER wildtype (*odr-10::GFP + lin-15(+)*) and NER defective *ercc-1* mutants (*odr-10::GFP + lin-15(+); ercc-1(tm1981) I*). We observed that CisPt increases the fraction of degraded (i.e., outgrowths harbouring) AWA neurons only after treatment with high concentration of CisPt (i.e. 300 µg/ml) for an extended period of time (i.e. 72 h) **(Fig. 1F, 1G)**. Importantly, such neurodegenerative processes were not observed using CisPt concentrations (i.e., 50 µg/ml) that already caused significant sensory dysfunction **(see Fig. 1A)**. As an additional control for DNA damage-induced injury, we employed UV-C, which induces bulky DNA lesions that are subject to NER repair as well. Using UV-C doses of 10-40 J/m^2^ that heavily impaired sensory function in the chemotaxis assay **(Supplementary Fig. S1C)**, we found a higher degree of morphological degradation in form of bead-like outgrowths as well as neuronal cell death, as indicated by fragmentation of the neuronal process in *ercc-1* mutants **(Supplementary Fig. S1A** and **S1B)** than after CisPt. These findings indicate, that CisPt concentrations leading to sensory dysfunction induce age-associated functional and transcriptional alterations rather than morphological abnormalities in the sensory AWA neurons.

### 3.2. Cholinergic and GABAergic signaling modulate CisPt-induced neuronal dysfunction

To investigate whether CisPt-mediated sensory dysfunction of ERCC-1 defective mutants is related to alterations in neurotransmitter levels, the amount of the neurotransmitters dopamine, serotonin, GABA and acetylcholine were quantitatively analyzed under basal situation and following 72 h of CisPt treatment using mass spectrometry-based method. Following treatment with a high CisPt concentration (i.e., 300 µg/ml), a significant increase in dopamine level was found, which is independent of the NER status **(Fig. 2A).** Most important, supplementation with dopamine did not protect *ercc-1* mutants from CisPt induced sensory dysfunction **(Fig. 2B)**, indicating that dopaminergic neurotransmission is not of functional relevance under our experimental setting. Also for serotonin, no significant alterations were found under basal situation and following CisPt treatment **(Fig. 2C)**. Moreover, serotonin supplementation did not ameliorate CisPt-induced sensory neurotoxicity **(Fig. 2D)**. Hence, we conclude that serotoninergic neurotransmission is also not of relevance in the context of CisPt-triggered sensory neurotoxicity in NER deficient nematodes. Glutamate supplementation did also not ameliorate CisPt-induced neurotoxicity as observed in the chemotaxis assay **(Supplementary Fig. S2)**. Analyzing GABA levels, we found a significant drop in both wildtype and *ercc-1* deficient nematodes following treatment with low CisPt concentration **(Fig. 2E)**, whereas *ercc-1* mutants revealed an increase only at very high CisPt concentration (i.e., 300 µg/ml) **(Fig. 2E).** Most important, supplementation with GABA protected the *ercc-1* mutants from CisPt-induced sensory dysfunction **(Fig. 2F)**. Based on this finding we conclude that a reduced GABAergic neurotransmission in CisPt treated *ercc-1* mutants is of relevance for their sensory dysfunction and, furthermore, that GABA acts in a neuroprotective manner. Acetylcholine levels remained unaltered in both strains following CisPt treatment **(Fig. 2G).** Noteworthy yet, supplementation with the acetylcholine agonist carbachol further aggraviated CisPt-mediated sensory neurotoxicity in *ercc-1* mutants **(Fig. 2H)**. Notably, carbachol mono-treatment already caused neurotoxic effects, as indicated by a drop in CI, in both wildtype and *ercc-1* mutated animals **(Fig. 2H)**, which, however, was stronger in *ercc-1* mutants **(Supplementary Fig. S3A)**. Most important, CisPt-induced sensory neurotoxicity was further enhanced by co-treatment with carbachol **(Fig. 2H)**. At the concentration used, carbachol had only moderate inhibitory effects on the nematodeś swimming behavior **(Supplementary Fig. S3B)**, indicating that it preferentially affects sensory functions of the nematode rather than motor functions.

**Fig. 2:**
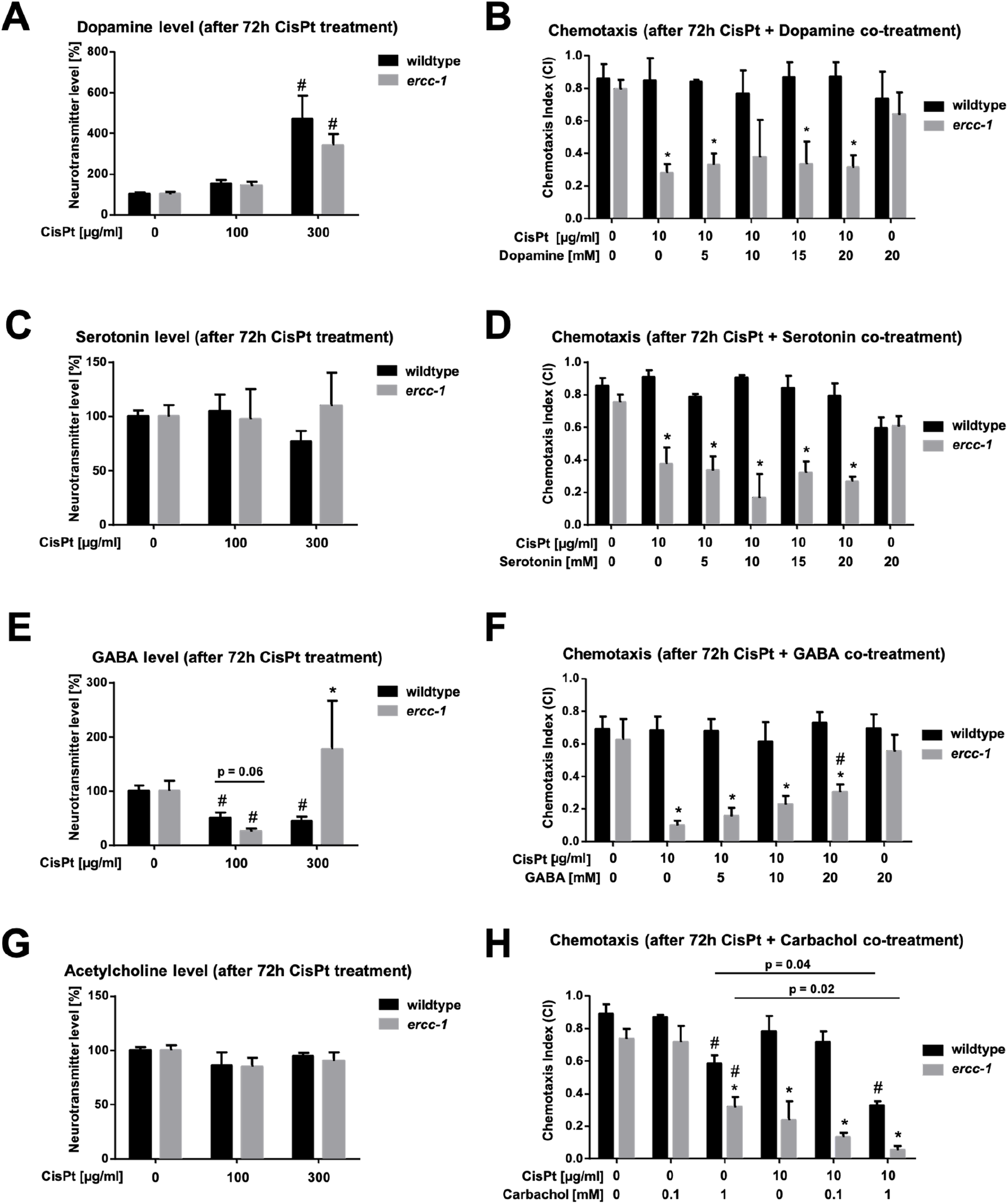
GABA signaling contributes to CisPt-induced sensory neurotoxicity. **A, C, E, G:** Mass spectrometry-based analysis of neurotransmitter levels of dopamine **(A)**, serotonin **(C)**, GABA **(E)** and acetylcholine **(G)** after 72 h of CisPt treatment (0-300 µg/ml) of wildtype and *ercc-1* animals. Data shown are the mean + SEM of two independent experiments with 4-5 biological replicates per experiment. *, p ≤ 0.05 (wildtype vs. mutant, students t-test); ^#^, p ≤ 0.05 (control vs. CisPt-treatment, One-Way ANOVA). **B, D, F, H:** Complementary chemotaxis analysis of wildtype and *ercc-1* mutated animals after 72 h of co-treatment with CisPt (10 µg/ml) and the respective neurotransmitters dopamine **(B)**, serotonin **(D)**, GABA **(F)** or carbachol **(H)** (concentrations as indicated). Data shown are the mean + SEM of three independent experiments (N ≥ 100 nematodes per experimental condition and experiment). *, p ≤ 0.05 (wildtype vs. mutant, students t-test); ^#^, p ≤ 0.05 (control vs. CisPt-treatment, One-Way ANOVA).

### 3.3. Opposing roles of GABAergic and cholinergic signaling indicate an excitation/inhibiton imbalance following CisPt exposure

For further clarification of the neuroprotective outcome of GABAergic signaling in the context of CisPt-mediated DNA damage-triggered sensory dysfunction, the CisPt response of different *C. elegans* mutants was investigated. Data obtained reveal that mutations in both GABA synthesis (*unc-25*) and GABA_A_ transmission (*unc-49*) impair basal sensory functionality **(Fig. 3A)**. Following CisPt treatment, only the *unc-49* mutant responded with a further decrease in CI **(Fig. 3A)**, which was not observed in the *unc-25* mutant **(Fig. 3A)**. Notably, *unc-49* mutated animals displayed a reduced basal swimming activity that was further impaired upon CisPt treatment **(Supplementary Fig. S4A)**. Notably, *unc-49* mutants were more susceptible to sensory neuronal dysfunction at lower CisPt concentrations in the chemotaxis assay (i.e. 10 µg/ml) than to motor defects monitored in the swimming assay (i.e. 50 µg/ml) **(Fig. 3A** and **Supplementary Fig. S4)**. Other mutants defective in GABA_B_ transmission (*gbb-1;gbb-2*) and ligand-gated chloride channels (*lgc-36, lgc-37*) neither revealed any differences in basal CI nor following CisPt exposure **(Fig. 3A)**, indicating that intact GABA_A_ signaling rather than GABA_B_ signaling is required for normal sensory function in *C. elegans*. Moreover, our data indicate that ionotropic GABA_A_ receptor signaling contributes to protection against CisPt-induced sensory neurotoxicity, while metabotropic GABA_B_ signaling appears to play a minor role.

**Fig. 3:**
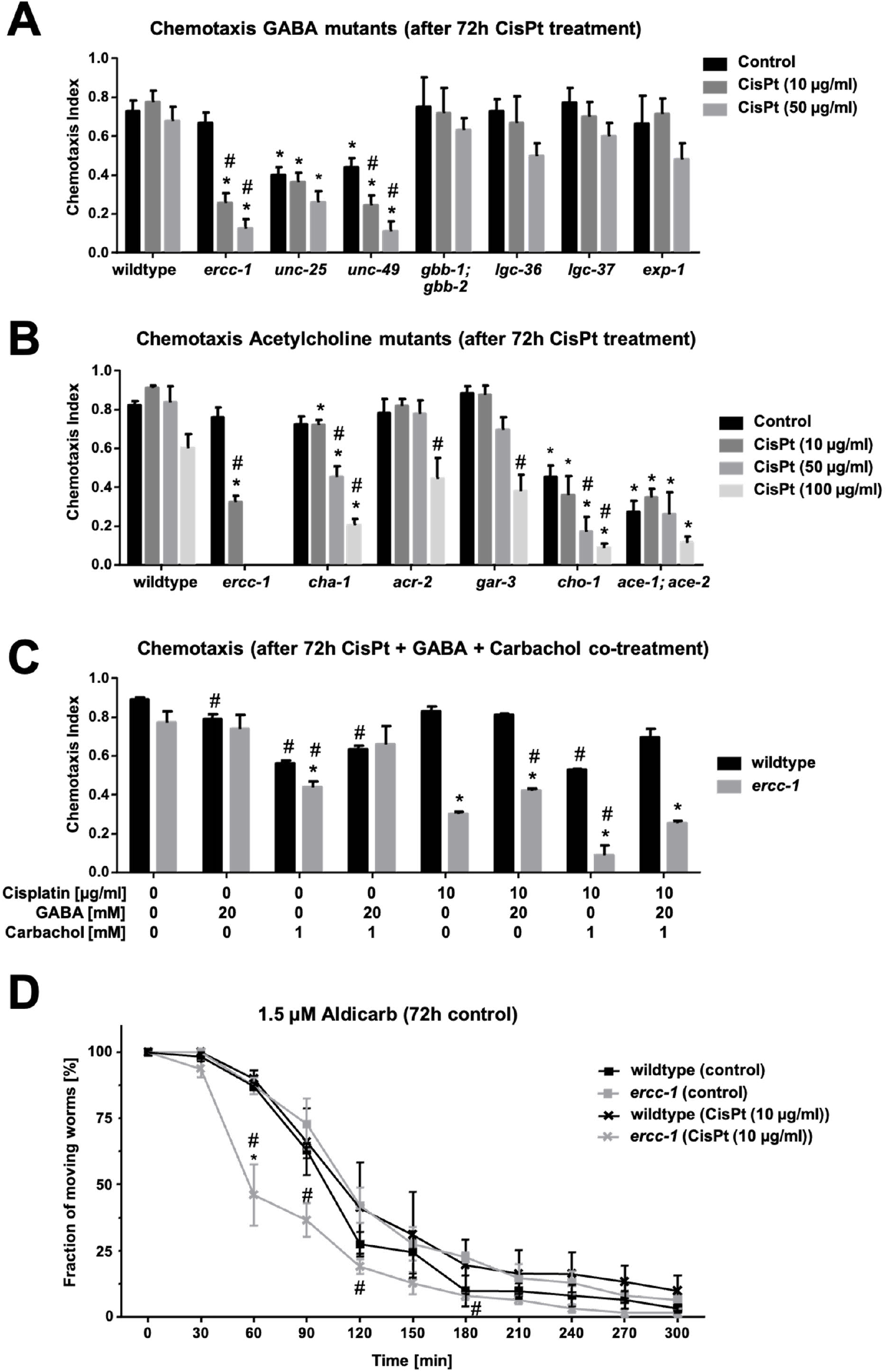
Both GABAergic and cholinergig signaling influence the vulnerability of *ercc-1* mutants to CisPt-induced neuronal dysfunction. **A:** Chemotaxis analysis of wildype and mutants of GABAergic signaling (*unc-25, unc-49, gbb-1;gbb-2, Igc-36, Igc-37, exp-1*) after 72 h of CisPt treatment (0-50 µg/ml). Data shown are the mean + SEM of three independent experiments (N ≥ 100 nematodes per experimental condition and experiment). *, p ≤ 0.05 (wildtype vs. mutant, students t-test); ^#^, p ≤ 0.05 (control vs. CisPt-treatment, One-Way ANOVA). **B:** Chemotaxis analysis of wildtype and various mutants of cholinergic signaling (*cha-1, acr-2, gar-3, cho-1, ace-1;ace-2*) after 72 h of CisPt treatment (0-100 µg/ml). Data shown are the mean + SEM of three independent experiments (N ≥ 100 nematodes per experimental condition and experiment). *, p ≤ 0.05 (wildtype vs. mutant, students t-test); ^#^, p ≤ 0.05 (control vs. CisPt-treatment, One-Way ANOVA). **C:** Chemotaxis analysis of wildtype and NER-defective *ercc-1* mutant after combined treatment with CisPt (0-10 µg/ml) ± carbachol (1 mM) ± GABA (20 mM) for 72 h. Data shown are the mean + SEM of three independent experiments (N ≥ 100 nematodes per experimental condition and experiment). *, p ≤ 0.05 (wildtype vs. mutant, students t-test); ^#^, p ≤ 0.05 (control vs. CisPt-treatment, One-Way ANOVA). **D:** Aldicarb assay of wildtype and *ercc-1* mutant animals after 72 h of CisPt treatment (0, 10 µg/ml) (N ≥ 60, n = 3). Data shown are mean ± SEM of three independent experiments with 18-21 nematodes being analyzed per experimental condition). *, p ≤ 0.05 (wildtype vs. mutant, students t-test); ^#^, p ≤ 0.05 (control vs. CisPt-treatment, One-Way ANOVA).

In view of the opposite effects observed upon co-treatment of CisPt with either GABA or carbachol, we additionally investigated the CisPt response of mutants defective in cholinergic signaling. In contrast to the effect observed in GABAergic mutants, neither the metabotropic receptor mutant *gar-3* nor the ionotropic receptor mutant *acr-2* revealed different responses to CisPt as compared to the wildtype **(Fig. 3B)**, while choline acetyltransferase mutant *cha-1* displayed reduced chemotaxis activity upon CisPt treatment (i.e., 50 µg/ml) **(Fig. 3B)**. Interestingly, choline transporter mutant *cho-1* showed an already reduced basal chemotaxis towards diacetyl, which was further reduced following CisPt exposure (i.e., 50 µg/ml) **(Fig. 3B)**. Similarly, acetylcholine esterase double mutant *ace-1;ace-2* also showed impaired chemotaxis under basal situation **(Fig. 3B)**. However CisPt treatment did not cause further impairment in its chemotaxis activity **(Fig. 3B)**. Since both GABAergic and cholinergic signaling appeared to be required for adequate sensory function, we next investigated the outcome of a triple co-treatment with CisPt, GABA and carbachol. The data obtained show that exogenous supplementation with GABA significantly ameliorated the neurotoxic effects following both carbachol and CisPt mono-treatment as well as combined treatment **(Fig. 3C).** This data indicate that GABA supplementation partially counteracts neuronal dysfunction induced by carbachol-mediated cholinergic overstimulation that is aggravated by additional CisPt exposure. In view of the reduced basal chemotaxis activity of *ace-1;ace-2*, we analyzed the influence of CisPt on acetylcholine esterase inhibitor aldicarb-induced paralysis of wildtype and *ercc-1* mutants. The data obtained show a similar time-dependent decrease in the nematodeś motility independent of the their NER status under basal conditions **(Fig. 3D).** By contrast, aldicarb-induced loss in motility was accelerated following CisPt treatment of *ercc-1* mutants **(Fig. 3D).** Moreover, to further scrutinize the relevance of GABAergic signaling in aldicarb-induced paralysis and its role in CisPt-induced neuronal dysfunction, we additionally included the *unc-49* mutant in our analysis. Strikingly, the *unc-49* mutant exhibited markedly increased sentitivity to aldicarb-induced paralysis as compared to the wildtype that was further exacerbated by CisPt-treatment **(Supplementary Fig. S5)**. Given that aldicarb-induced paralysis is promoted by an imbalance between excitatory and inhibitory signaling, this finding further suggests that CisPt-induced DNA damage shifts the balance between cholinergic excitation and GABAergic inhibition towards an excessive excitation, either through enhanced cholinergic excitation, reduced GABAergic inhibiton or both.

### 3.4. GABA supplementation reduces CisPt-induced degeneration of GABAergic neurons

To further elucidate the mechanisms of CisPt-mediated reduction in GABA levels and reduced chemotaxis, we investigated the influence of CisPt on the morphological integrity of GABAergic neurons. We found that CisPt substantially increases beading of GABAergic neurons in both NER wildtype (*unc-47p::GFP*) and NER-defective *ercc-1* (*unc-47p::GFP; ercc-1*) mutants **(Fig. 4A**, **Fig. 4B)**. In contrast to the observed neuronal degradation in AWA neurons, we found an increased rate of outgrowths on GABAergic neurons already at low concentrations of CisPt (i.e., 10 µg/ml) exclusively in *ercc-1* mutants **(Fig. 4A**, **Fig. 4B)**. Additionally, high concentration of CisPt (i.e., 100 µg/ml) evoked cell death of GABAergic neurons in *ercc-1* mutants, as visible by fragmentation of the neuronal process **(Fig. 4B**, indicated by arrows). Notably, this CisPt-stimulated cytotoxic response can be prevented by supplementation with GABA **(Fig. 4A**, **Fig. 4B)**. Based on this finding we hypothesize that CisPt-induced DNA damage-mediated sensory dysfunction involves the degeneration of GABAergic neurons, leading to a drop in GABA levels.

**Fig. 4:**
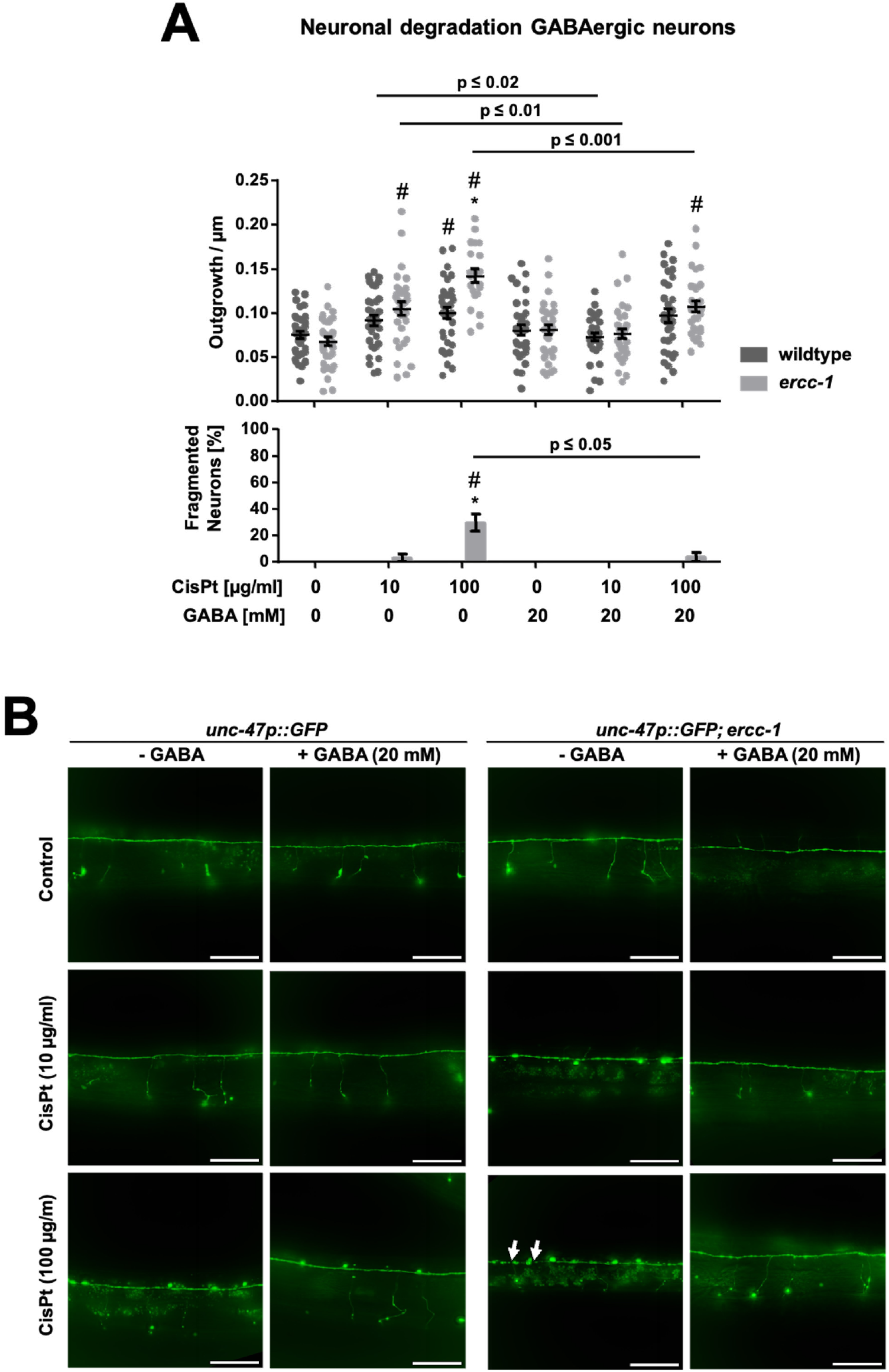
GABA substitution rescues degradation of GABAergic neurons. **A:** Quantification of neuronal degradation of GABAergic neurons in NER-wildtype and *ercc-1* mutated nematodes after combined treatment with CisPt (0-100 µg/ml) ± GABA (20 mM) for 72 h with representative images **(B)**. Scale bar = 50µm. Arrows indicate interruptions of the neuronal process interpreted as fragmentation/ neuronal cell death. Quantitative data shown are from n = 3 independent experiments, with at least 10 images being analyzed per experimental condition.*, p ≤ 0.05 (wildtype vs. mutant, students t-test); ^#^, p ≤ 0.05 (control vs. CisPt-treatment, One-Way ANOVA).

### 3.4. NER deficiency enhances the transcriptional response to cisplatin and dysregulates neurotransmission-associated genes

For a more in depth comparative analysis of the molecular stress responses evoked by CisPt and their putative relevance for the observed preferential sensory dysfunction in NER defective nematodes, alterations in the transcriptome were analyzed under basal situation and following CisPt treatment of wildtype and *ercc-1* mutated worms by RNA-seq-based analysis The differentially expressed genes were subjected to functional annotation clustering using the DAVID bioinformatics platform to identify biological processes and signaling pathways that are enriched in CisPt treated *ercc-1* mutants.

To identify genotype-dependent differences independent of CisPt treatment, we first compared the transcriptome of untreated wildtype versus *ercc-1* mutated animals. A total of 940 genes were upregulated and 147 genes were downregulated in *ercc-1* mutated animals as compared to the wildtype under basal conditions **(Fig. 5A)**. Functional annotation clustering revealed a prominent enrichment of genes associated with “Protein signaling and ATP dependent processes” and “GABA transport”. In contrast, downregulated genes were primarily associated with “Transcriptional regulation and nuclear receptor signaling” as well as “Stress response and longevity pathways” **(Fig. 5B)**. Comparison of CisPt-treated wildtype versus CisPt-treated *ercc-1* mutants revealed a total of 1296 upregulated and 49 downregulated genes in the NER-defective mutant as compared to the wildtype **(Fig. 5C)**. These differentially expressed genes were largely enriched in the same biological processes as under basal conditions **(Fig. 5D)**.

**Fig. 5:**
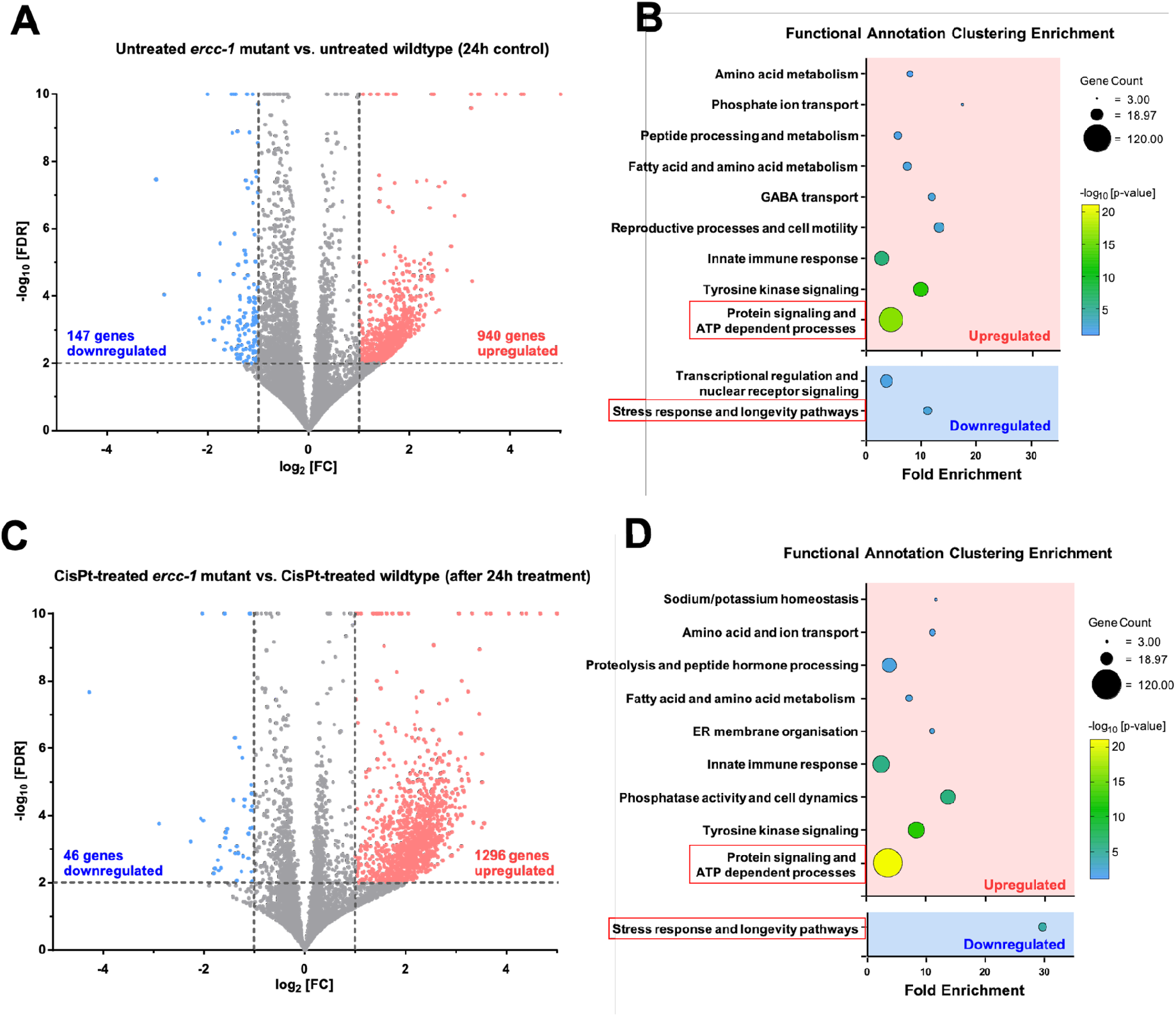
CisPt broadens ERCC-1 deficiency-associated transcriptional responses. RNA-seq analysis of wildtype and *ercc-1* animals under basal situation and 24 h after CisPt treatment (100 µg/ml) (N = 4 biological replicates per experimental condition). **A:** Volcano plot comparison of genes expressed in untreated wildtype vs. untreated *ercc-1* mutated *C. elegans*. Upregulated genes (-log_10_(FDR) ≥ 2) were shown in red (log_2_[FC] ≥ 1), downregulated genes in blue (log_2_[FC] ≥ -1). For visualization, values were capped at - log_10_[FDR] = 10 and log_2_[FC] = ±5. **B:** Corresponding functional enrichment annotation clustering plot of differentially expressed genes in untreated wildtype vs. untreated *ercc-1* mutated *C. elegans*. Red boxes indicate selected clusters for in depth analysis (see **Supplementary Fig S6)**. **C:** Volcano plot comparison of differentially expressed genes in treated wildtype vs. treated *ercc-1* mutated *C. elegans.* Upregulated genes (-log_10_(FDR) ≥ 2) were shown in red (log_2_[FC] ≥ 1), downregulated genes in blue (log_2_[FC] ≥ -1). For visualization, values were capped at - log_10_[FDR] = 10 and log_2_[FC] = ±5. **D:** Corresponding functional enrichment annotation clustering plot of differentially expressed genes in CisPt treated wildtype vs. CisPt treated *ercc-1* mutated *C. elegans*. Red boxes indicate the clusters that were selected for further in depth analysis (see **Supplementary Fig S6)**.

Because of some similarities of functional enrichment clusters between untreated or CisPt treated wildtype and *ercc-1* mutants, we performed a more in depth analysis of the genes represented in selected shared clusters, which we speculate to be relevant for the observed neuronal impairment following CisPt treatment. The upregulated cluster “Protein signaling and ATP dependent processes” appeared in both comparative analyses **(Fig. 5B** and **Fig. 5D)**. Analysing specifically upregulated genes that are represented in this shared cluster, a total of 90 genes were identified that were specifically upregulated in *ercc-1* mutants as compared to wildtype animals under basal condition **(Supplementary Fig S6A)**, whereas 115 genes were selectively upregulated in the NER mutants following CisPt exposure **(Supplementary Fig S6B)**. In both groups there was a substantial overlap of 77 differentially expressed genes that are involved in protein kinase signaling (e.g., *kin-5, kin-21, kin-24*), ATP-dependent transport (e.g., *abt-1, catp-4, wht-8*), insulin-like signaling (*ins-31*) and neuronal protein phosphorylation (e.g., *ttbk-2.1, ttbk-5, ttbk-8.2*). Specifically after CisPt exposure, another 38 genes were additionally represented in the cluster, widening the pre-existing transcriptional signature to genes related to ATP-dependent transport (e.g., *abt-5, catp-3, wht-4, wht-5*), insulin-like signaling (*ins-37*), DNA-repair (*rad-50)* and neuronal protein phosphorylation (e.g., *ttbk-2.2, ttbk-3, ttbk-4, ttbk-8.1*). This finding demonstrates a further sophisticated shaping of the transcriptional response upon genotoxic insult caused by CisPt under situation of ERCC-1 deficiency. On the other hand, the downregulated cluster “Stress response and longevity pathways” was also represented in both untreated or CisPt treated wildtype and *ercc-1* mutants **(Fig. 5B** and **Fig. 5D)**. Here, 9 genes were downregulated under basal situation in *ercc-1* mutants as compared to wildtype animals **(Supplementary Fig S6C)** and 8 genes after CisPt treatment **(Supplementary Fig S6D)**. The expression of five heatshock proteins (i.e., *hsp-16.41*, *hsp-16.48*, *hsp-16.49*, *hsp-70* and *rskn-2* (homologue of mammalian RSK)) was altered in *ercc-1* mutants as compared to the wildtype, both under basal situation and following CisPt treatment. However, the expression of the stress regulating genes *kpc-1*, *fipr-24*, *lea-1* and *jmjc-1* was exclusively represented under basal conditions. Following CisPt treatment, the cluster included additional heatshock proteins (i.e., *hsp-16.2*, *hsp-16.20*) and *C16E9.1*, indicating that the ERCC-1 deficiency-associated transcriptional signature is characterized by a CisPt-mediated suppression of heatshock proteins that are linked to aging and longevity.

To assess genotype-specific transcriptional responses to CisPt treatment as related to the NER repair status of the nematodes, we next compared CisPt-treated wildtype and *ercc-1* mutated animals with their respective untreated control. CisPt treatment induced marked transcriptional changes in both wildtype an *ercc-1* mutants. In wildtype animals, 655 genes were upregulated and 78 genes were downregulated following CisPt exposure **(Fig. 6A),** with functional annotation analysis revealing enrichment particularly in “Neuronal signaling” **(Fig. 6B)**. The genes represented in this cluster comprise 64 genes involved in various different neuronal functions **(Supplementary Fig. S7A),** including cholinergic (e.g., *unc-17, acr-19, unc-38, lev-1, acc-3*), glutamatergic (e.g., *glr-5, glc-3, glc-4, avr-14*) and GABAergic (i.e. *snf-11, gab-1, gbb-2*) neurotransmission, whereas dopaminergic neurotransmission was only represented by one gene (*dat-1*). Another prominently enriched cluster was “GPCR signaling” **(Fig. 6A)**. The 47 genes represented in this cluster **(Supplementary Fig. S7B)** are prominently neuropeptide G protein-coupled receptor (GPCR) genes from the *npr* and *dmsr* family, adenylyl cyclases (*acy-1* and *acy-2*), insulin-like peptides (*ins-1* and *ins-20*) as well as downstream components *gpa-2* and *gpa-11,* which signal via cAMP and the DAF-16 FOXO transcription factor family, respectively, as well as the downstream effector *odr-3* of the chemosensory GPCR *odr-10*. Interestingly, the prominent enrichment of GPCR-related genes was observed exclusively in wildtype animals, demonstrating large deficits of *ercc-1* mutants regarding CisPt-stimulated activation of GPCR-regulated signaling.

**Fig. 6:**
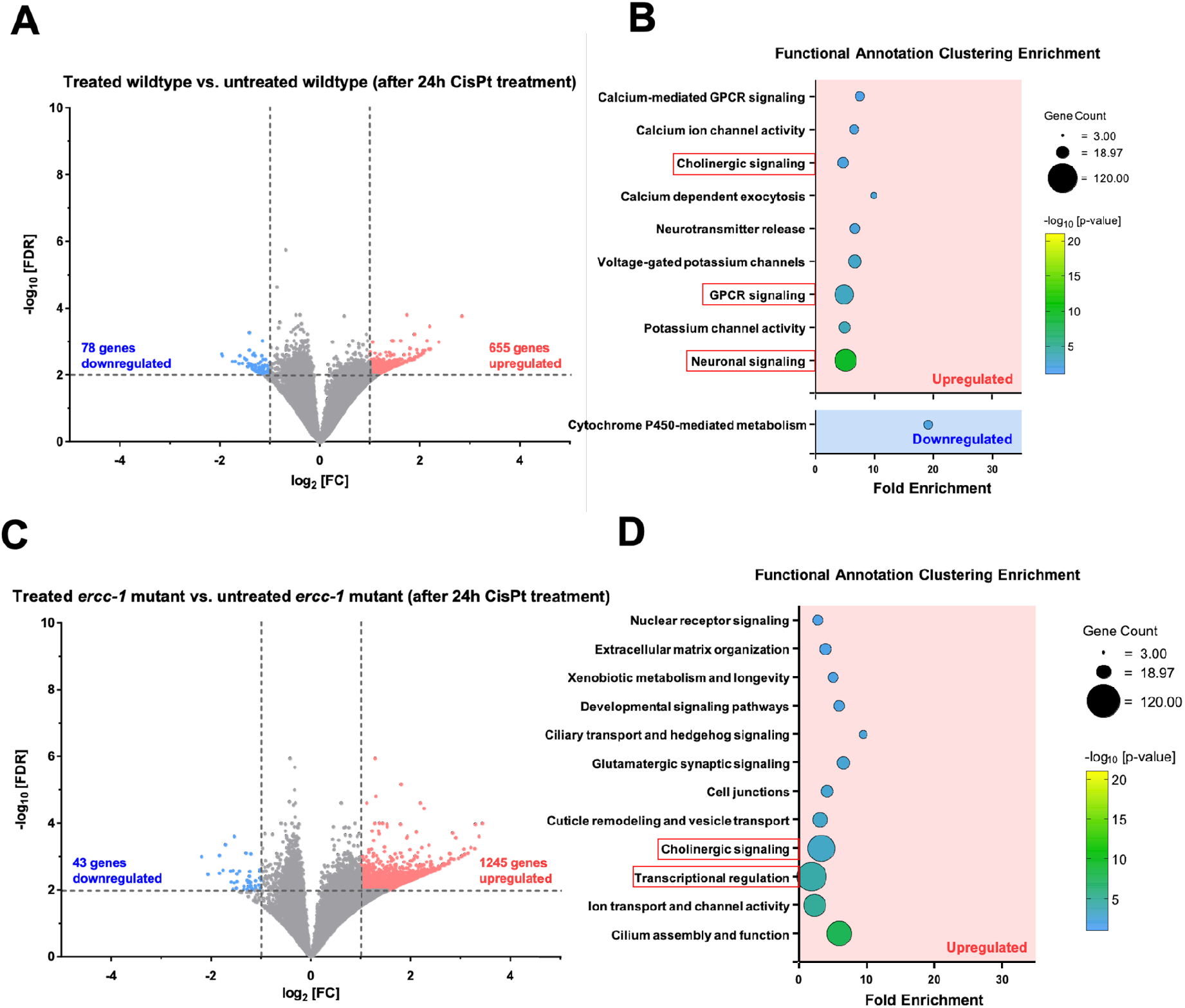
Wildtype nematodes and *ercc-1* mutants exhibit a distinct transcriptional response to CisPt. RNA-seq analysis of wildtype nematodes and *ercc-1* mutants was performed 24 h after CisPt treatment (100 µg/ml) (N = 4 biological replicates per experimental condition). **A:** Volcano plot comparison of expressed genes in untreated vs. CisPt treated wildtype *C. elegans.* Relevant genes (-log_10_(FDR) ≥ 2) were shown in red when upregulated (log_2_[FC] ≥ 1) or blue when downregulated (log_2_[FC] ≥ -1). For visualization, values were capped at - log_10_[FDR] = 10 and log_2_[FC] = ±5. **B:** Corresponding functional enrichment annotation clustering plot of differentially expressed genes in untreated vs. CisPt treated wildtype *C. elegans*. Red boxes indicate selected clusters for in depth analysis (see **Supplementary Fig. S7** and **Supplementary Fig. S8)**. **C:** Volcano plot comparison of expressed genes in untreated vs. CisPt treated *ercc-1* mutated *C. elegans*. Relevant genes (-log_10_(FDR) ≥ 2) were shown in red when upregulated (log_2_[FC] ≥ 1) or blue when downregulated (log_2_[FC] ≥ -1). For visualization, values were capped at - log_10_[FDR] = 10 and log_2_[FC] = ±5. **D:** Corresponding functional enrichment annotation clustering plot of differentially expressed genes in untreated vs. CisPt treated *ercc-1* mutated *C. elegans*. Red boxes indicate selected clusters for in depth analysis (see **Supplementary Fig. S7** and **Supplementary Fig. S8)**.

A substantially stronger transcriptional response to CisPt treatment was observed in *ercc-1* mutant animals, with a considerably larger amount of 1245 upregulated and 43 downregulated genes **(Fig. 6C)**. Notably, the most upregulated genes were enriched in the cluster “Transcriptional regulation” **(Fig. 6D)**. This cluster is represented by 90 genes that are exclusively enriched in CisPt treated *ercc-1* mutant animals **(Supplementary Fig. S7C)**. We identified several transcriptional regulators, including 12 homeobox proteins (e.g., *ceh-2 ceh-6, ceh-17*), 24 nuclear hormone receptors with transcription activating functions (e.g., *nhr-18, nhr-25, nhr-52*) and several transcription factors involved in the regulation of neuronal identity and differentiation (e.g., *unc-3, unc-30, unc-86*). These results highlight a transcriptional remodeling of neuronal identity programs resulting from persisting CisPt-induced DNA damage in NER defective *ercc-1* mutants, which may contribute to their enhanced sensory dysfunction.

Furthermore, the cluster “Cholinergic transmission” was commonly upregulated in both CisPt-treated wildtype and *ercc-1* mutated animals **(Fig. 6B** and **Fig. 6D)**. Following CisPt exposure, the respective cluster contained 14 genes upregulated in the wildtype **(Supplementary Fig. S8A)** and 74 genes in the *ercc-1* mutant **(Supplementary Fig. S8B)**. Only three genes (i.e., *lgc-8, lgc-11, lgc-21*) were shared between wildtype and *ercc-1* mutant. Wildtype specifically upregulated genes were predominantly associated with acetylcholine transport (*unc-17*) and metabotropic and ionotropic receptors (e.g., *eat-2, acr-2, acr-12*). By contrast, *ercc-1* specific genes included different types of acetylcholine receptors (e.g., *gar-2, gar-3, acr-19*) as well as genes involved in neurotransmitter release (e.g., *nrx-1, nlg-1, rab-3*), neuronal excitability (i.e. *unc-2*) and mechanosensory signaling (i.e. *mec-2*, *mec-4*).

Collectively, the results from our transcriptome analysis demonstrate that CisPt treatment further emphasizes pre-existing differences in the transcriptomic signature between wildtype and *ercc-1* mutants, including neuronal kinase signaling and insulin-like signaling involved in aging and longevity. Moreover, genotype specific responses to CisPt comprise the upregulation of genes involved in neuronal communication and GPCR signaling in the wildtype, whereas the *ercc-1* mutated animals are characterized by upregulation of genes that are predominantly involved in the regulation of cholinergic transmission and transcriptional regulation of neuronal identity and differentiation programs.

## 4. Discussion

Platinating anticancer drugs cause severe and long-lasting adverse neurotoxic effects, including cognitive impairment and CIPN that affect the patients’ quality of life (55, 56). The impact of platinating agent-induced DNA damage in the development of sensory neurotoxicity is largely unknown. In order to elucidate the contribution of insufficient NER-mediated repair of CisPt-induced DNA damage in CisPt-mediated sensory neurotoxicity, we employed the nematode *C. elegans* as a powerful 3R-conform model in basic toxicological research (32). Previously we investigated the outcome of a NER defect caused by ERCC-1 deficiency on CisPt-mediated sensory functions of *C. elegans,* demonstrating that CisPt-induced DNA crosslinks distort the sensory ability of the nematode (43). We showed that CisPt treatment impairs chemotaxis at much lower concentration than swimming activity in NER defective *ercc-1* mutants, indicating a preferential DNA damage-related toxicity on sensory neurons over motor neurons as suggested before (43). This finding is in line with data demonstrating that CisPt-induced damage to skeletal muscles majorly involves DNA damage independent mechanisms (57), which may require higher CisPt concentrations and, furthermore, that the sensitivity of neurons to DNA damage is substantially higher than that of muscle cells (42).

The main aim of the present study was to further elucidate the molecular mechanisms of CisPt-induced DNA damage-related sensory toxicity. Noteworthy, while the NER-defective *ercc-1* mutants revealed a large hypersensitivity to CisPt-induced sensory neurotoxicity, such effects were not observed in mutants of the DDR-related factors p53, ATR and ATM. This finding indicates that lack of ERCC1-mediated repair of CisPt-induced DNA damage, rather than the subsequent activation of ATM/ATR- and p53-related mechanisms of the DDR are of relevance for sensory neurotoxicty evoked by CisPt. Hence, we speculate that CisPt-induced primary DNA lesions or secondary DNA lesions resulting from their insufficient or error-prone processing are ultimately triggering sensory neurotoxicity. Since ERCC-1 is part of both the GG-NER and TC-NER machinery, it appears feasible that transcriptional deficits of CisPt-treated *ercc-1* mutants are of relevance for their hypersensitive phenotype. This hypothesis gains support by the fact that mutants lacking the TC-NER-related factor CSB also reveal enhanced sensitivity to CisPt-induced sensory neurotoxicity, whereas mutation of the GG-NER-related factor XPC has only minor effect (43). Moreover, our hypothesis is also in line with the finding that CSB-related functions are involved in DNA damage-triggered neurodegeneration and aging of the nematode (9) and, moreover, sensory functions decline with aging (58). Thus, it is rational to assume that the severe CisPt-mediated sensory dysfunction observed in *ercc-1* mutants is due to DNA damage-induced accelerated aging processes driven by insufficient TC-NER-related processes. This hypothesis gains further support by the fact that premature aging can result from loss of ERCC-1 in NER-defective *C. elegans* mutants (59) as well as in *ercc-1* deficient mice (60-62). Further discussing the influence of NER on platin-induced neurotoxicity, it is noteworthy that gene variants of the NER-related genes ERCC-4 and ERCC-6 affect peripheral neuropathy caused by oxaliplatin in humans (63). Moreover, ERCC-1 polymorphism increases the risk of platinating agent-induced nephrotoxicity (64), which is another clinically relevant adverse effect of platin-based anticancer therapy. Taken together, consistent data obtained across different species strongly indicate that ERCC1-mediated TC-NER of CisPt-induced DNA lesions mitigates multiple dose-limiting adverse effects of platinating agents, including neuro- and nephrotoxicity.

Since it is rational to assume that neurotoxicity following CisPt exposure of *ercc-1* mutants involves alterations in mechanisms related to neurotransmission (15), we next analyzed the levels of various candidate neurotransmitters. We found that CisPt treatment selectively reduces GABA neurotransmitter levels in both wildtype and *ercc-1* mutated animals while the amount of other neurotransmitters remained largely unaffected by drug treatment. Previously we demonstrated that the basal mRNA expression of genes coding for factors regulating neurotransmitter synthesis and neurotransmission by acetylcholine, dopamine, glutamate, serotonin and GABA is downregulated in *ercc-1* mutants (43). Thus, mRNA expression data do not fully reflect alterations in neurotransmitter levels as analyzed by mass spectrometry-based method. Taken together, we hypothesize that disturbance of GABAergic neurotransmission following CisPt treatment is of utmost relevance for DNA damage driven sensory neurotoxicity observed in the absence of ERCC-1. This interpretation is further supported by the observed CisPt-induced sensory defect in GABA_A_ signaling mutants (i.e., *unc-49*) as detected in the chemotaxis assay. Noteworthy, CisPt-induced neurotoxicity, as analyzed on the level of pharyngeal pumping activity, mainly involves dopaminergic and serotonergic neurotransmission (37, 39), which is however not of major relvance for sensory neurotoxicity evoked by the platinating agent as demonstrated in the present study. Thus, different neurotransmitter systems contribute to CisPt-induced neurotoxicity, with disturbed GABAergic neurotransmission being a key neurotoxic response resulting from CisPt-mediated damage to chemosensory neurons. In line with our data, imbalances in GABAergic signaling also contribute to CisPt-induced neuropathy in rats (65). In addition to GABA-related mechanisms, we observed cholinergic signaling to further promote CisPt-induced sensory dysfunction under situation of NER deficiency. Conversely, mutants of choline acetyltransferase (i.e., *cha-1*) and choline transport (i.e., *cho-1*) were sensitive to CisPt treatment in the chemotaxis assay. This is in line with recent findings linking neuronal NER activity to acetylcholine signaling in promoting systemic resistance to DNA damage (66). We suggest that, while both GABAergic and cholinergic neurotransmitter systems are apparently required for normal sensory function, disruption of the balance between GABAergic inhibition and cholinergic excitation might contribute to CisPt-induced sensory dysfunction as a promoting factor. This hypothesis gains support by our observed GABA supplementation-mediated alleviation of the detrimental effects resulting from a combined carbachol and CisPt treatment in the chemotaxis assay. Furthermore, CisPt-treated *ercc-1* mutants were hypersensitive to aldicarb-induced paralysis, a phenotype that has been associated with alterations in synaptic transmission (53). In this context, it is noteworthy that cholinergic factors also interfere with developmental neurotoxicity evoked by the heavy metal nickel (67), supporting the hypothesis that the relevant neurotransmitters involved in agent-induced neurotoxicity depend on the specific kind of neurotoxic stimulus. In line with this, numerous mechanisms have been identified to contribute to platinating agent-induced CIPN in humans (15, 68).

Another important question is whether CisPt-induced sensory dysfunction related to GABAergic neurotransmission is due to neuronal cells death. Our previous findings indicated that sensory impairment occurrs in the absence of detectable apoptotic nor necrotic loss of AWA neurons, suggesting that neuronal dysfunction may precede cell death of the primary chemosensory neurons (43). In the present study we observed that CisPt treatment results in pronounced morphological alterations of GABAergic neurons in the form of a severe neuronal beading at low CisPt concentrations and even cell death at high concentrations of CisPt, particularly in *ercc-1* mutants. Consistent with the GABA-mediated rescue of the limited chemotaxis in CisPt-treated *ercc-1* mutants, GABA supplementation also attenuated neuronal beading, further supporting a neuroprotective role of GABA in the context of CisPt-induced neurotoxicity. Overall, this data show that enhanced sensory dysfunction of CisPt treated *ercc-1* mutants exposed to low concentrations of CisPt is likely not due to neuronal cell death but rather to an early functional and structural damage to GABAergic neurons. The obeserved CisPt-triggered beading process is suggested to represent a sign of progressive degeneration resembling morphological alterations that are frequently observed in aging, further supporting our hypothesis that DNA damage-induced accelerated aging is playing a major role in CisPt-induced sensory dysfunction. This interpretation is supported by other studies linking GABAergic signaling to neuronal aging in *C. elegans* (69-71). For example, loss of inhibitory GABAergic signaling and subsequent disruption of excitatory/inhibitory balance was previously described in aged worms (70). Similar to our results, this study demonstrated that pharmacological enhancement of ionotropic GABA_A_ signaling ameliorates neuronal aging while mutation of GABA_A_ receptor *unc-49* leads to a contrary aggravating aging effect (70). Additionally, GABA was also shown to confer direct neuroprotective effects through a *tmc-1* (transmembrane channel-like 1) dependent pathway promoting metabotropic GABA_B_ signaling, inhibiting the PKC pathway and, consequently, delaying structural and functional age-associated processes (71). Overall, these findings support the hypothesis that CisPt-induced impairment of GABA homeostasis contributes to neuronal dysfunction by further promoting features of neuronal aging and degeneration as well as a subsequential shift towards excitation caused by loss of inhibitory neuronal functions, especially under situation of persisting DNA damage in NER defective *ercc-1* mutants. Regarding other neuronal functions that become negatively affected by CisPt exposure, we previously showed that CisPt-mediated impairment of pharyngeal pumping is also independent of cell death of amphid-type neurons (37), further supporting the hypothesis that CisPt-mediated neurotoxicity is primarily based on an exacerbation of degeneration-associated neuronal dysfunction rather than neuronal cell death. Noteworthy is this context, CisPt induced neurotoxicity as evalulated by monitoring locomotion is independent of morphological alterations of dopaminergic neurons as well (39). This finding incidates that CisPt-mediated motor neurotoxicity is likely independent of dopaminergic mechanisms, which have recently been identified as relevant for memory-associated functions of the nematode (72).

Hypothesizing that the hypersensitivity of *ercc-1* mutants to CisPt-induced sensory neurotoxicity involves mechanisms of TC-NER, we comparatively analyzed the transcriptome of wildtype and *ercc-1* mutants, both under basal situation and following CisPt-treatment, in order to identify the molecular mechanism involved. Data obtained from extensive transcriptomic analyses indicate that CisPt-induced DNA damage in NER-defective *ercc-1* mutant animals further amplifies pre-existing differences in the mutant’s basal transcriptome as compared to the wildtype. These reinforced pathways in *ercc-1* mutants include neuronal kinase-related signaling mechanisms as well as longevity-related insulin-like signaling pathways. Wildtype animals predominantly upregulate the expression of genes that are related to a broard spectrum of neuronal communication mechanisms and neuronal GPCR signaling in response to CisPt treatment. Notaby, genes represented in the GPCR cluster prominently comprise neuropeptide GPCR genes from the *npr* and *dmsr* family, as well as downstream components like Gα-subunits *gpa-2* and *gpa-11* and adenylyl cyclases *acy-1* and *acy-2* involved in cyclic nucleotide-dependent signaling. These results point to post-synaptic receptor-mediated mechanisms that are impacted by CisPt under normal (i.e., wildtype) situation. This finding is of particular interest in the context of aging because, among other GPCR downstream components, *gpa-11* has been linked to DAF-16/FOXO-dependent regulation of longevity (73). Moreover, the CisPt-mediated upregulation of the insulin-like peptides *ins-1* and *ins-20* could indicate a link between CisPt-stimulated stress responses and insulin/IGF-1 signaling, which regulates stress resistance, longevity and the rate of aging (70, 74, 75). Most important, the prominent enrichment of GPCR-related genes was observed exclusively in wildtype animals, demonstrating large deficits of *ercc-1* mutants regarding CisPt-stimulated activation of GPCR-regulated signaling. This is of particular interest having in mind that GPCR are involved in CisPt-induced peripheral neuropathy in humans and are considered as targets for peripheral neuropathic pain (76, 77). Unlike wildtype nematodes, *ercc-1* mutated animals exhibited a pronounced CisPt-induced transcriptional activation of genes encoding transcription factors that are indicative of a remodeling of cholinergic transmission and activation of neuronal differentiation and identity. It remains unclear whether these CisPt-induced stress responses of *ercc-1* mutants contribute to their enhanced sensory dysfuntion or, alternatively, reflect the ERCC-1 mutantś attempt of a compensatory adaptive response. Moreover, *ercc-1* specific genes upregulated in response to CisPt include different types of acetylcholine receptors (e.g., *gar-2, gar-3, acr-19*) as well as of genes involved in neurotransmitter release (e.g., *nrx-1, nlg-1, rab-3*). This finding indicates that CisPt impacts both post-synaptic and pre-synaptic mechanisms related to neurotransmitter release in *ercc-1* mutants. Such mechanisms are also well-known to be affected in CIPN in humans (15).

## 5. Summary and Conclusion

In conclusion, our findings demonstrate that persisting DNA damage resulting from insufficient DNA repair of CisPt-induced DNA lesions promotes sensory dysfunction in NER-defective *ercc-1* mutated *C. elegans*. We hypothesize, that CisPt disrupts the physiological balance between GABAergic inhibition and cholinergic excitation, as indicated by reduced GABA levels and rescue of CisPt-mediated dysfunction by GABA supplementation as well as enhanced aldicarb sensitivity and elevated carbachol-induced sensory dysfunction following CisPt exposure. Moreover, these pharmacological data are reflected by genetic data obtained from the analysis of mutants of both GABAergic and cholinergic functions. Therefore, we hypothesize that balanced GABAergic and cholinergic mechanisms are required for healthy sensory functions and become disturbed by insufficient repair of CisPt-induced DNA damage in *ercc-1* mutants. Moreover, we show that *ercc-1* mutant animals display a particularly increased vulnerability of GABAergic neurons to CisPt-treatment as reflected by morphological alterations that are indicative of degenerative processes, which also explain the reduced GABA levels observed in the mutants as compared to the wildtype. On the transcriptomic level, unrepaired CisPt-induced DNA damage amplifies already pre-existing transcriptional signatures of longevity-related and neuronal kinase signaling-associated genes in *ercc-1* mutants. Furthermore, CisPt evokes distinct genotype-specific transcriptional responses that are indicative of a broard neuronal- and GPCR-related signaling exclusively in wildtype animals and activation of cholinergic signaling and neuronal identity programs specifically in *ercc-1* mutants. Taken together, these results suggest the loss of GABAergic integrity and the resulting shift towards cholinergic excitation as key mechanisms of CisPt-induced DNA damage-driven sensory neurotoxicity in NER defective *ercc-1* mutants **(Fig. 7, graphical abstract)**.

**Fig. 7:**
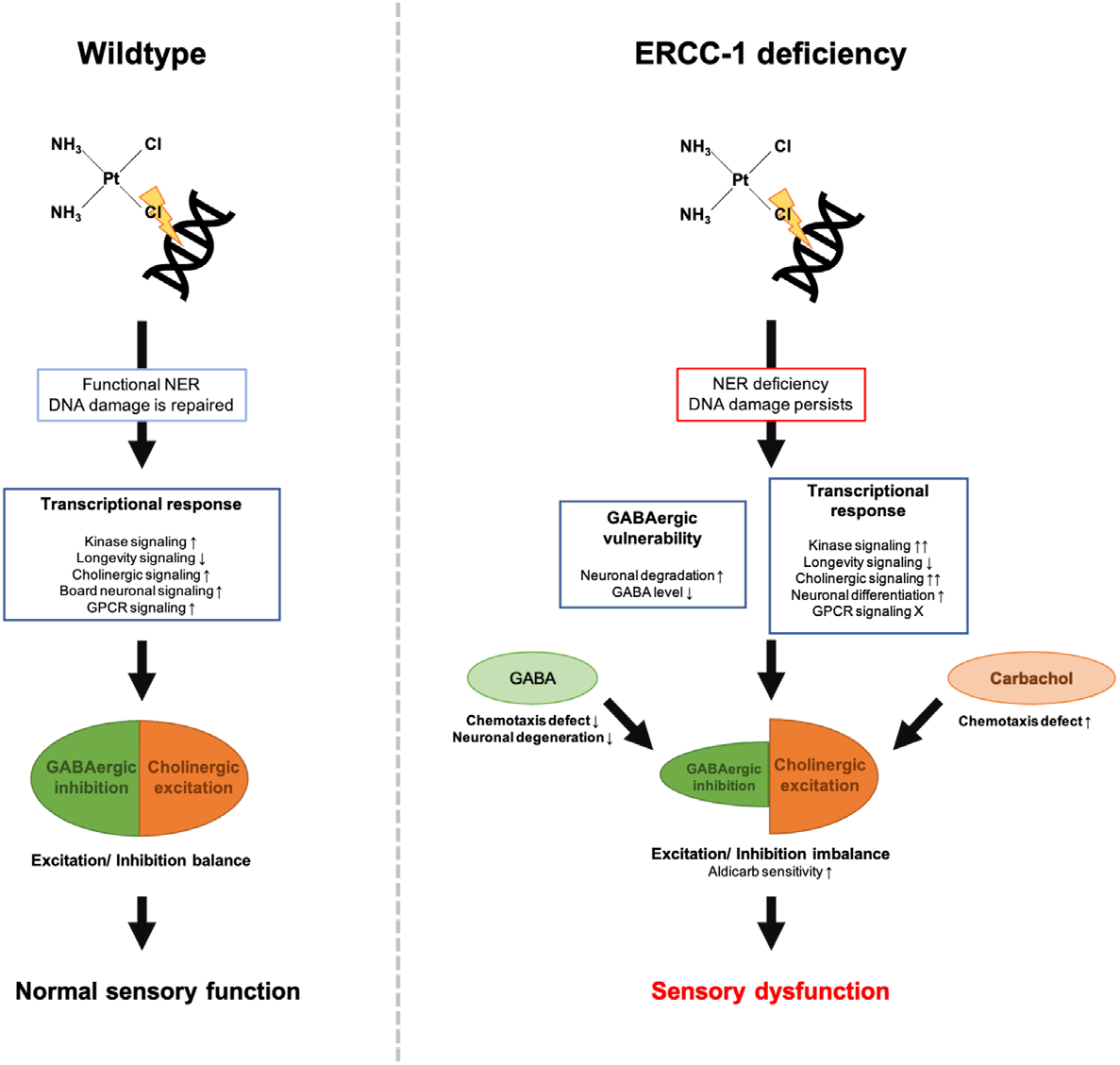
Proposed mechanisms of CisPt-induced sensory dysfunction in NER mutants of *C. elegans.* Insufficient DNA repair of CisPt-induced damage causes sensory dysfunction in NER defective *ercc-1* mutants. We propose increased GABAergic vulnerability, which is characterized by a pronounced neuronal degeneration in CisPt-treated *ercc-1* mutants and reduced GABA levels, to play a key role in the pathophysiology of CisPt-evoked sensory neurotoxicity. We hypothesize that the resulting loss of inhibitory signaling leads to a shift in the excitatory/inhibitory balance towards cholinergic excitation. This model is supported by a neuroprotective effect of GABA supplementation and carbachol-induced exacerbation of the CisPt-mediated sensory defect as well as increased aldicarb sensitivity of *ercc-1* mutants after CisPt treatment. Transcriptomic analysis further revealed distinct differences between wildtype and *ercc-1* mutants with broard neuronal- and GPCR-related signaling pathways being upregulated exclusively in CisPt treated wildtype nematodes, whereas *ercc-1* mutants are characterized by an amplification of cholinergic signaling- and neuronal identity-associated pathways in response to CisPt exposure.

## Acknowledgment

We thank the Caenorhabditis Genetics Center which is funded by the NIH Office of Research Infrastructure Programs (P40 OD010440), the *C. elegans* Gene Knockout Project at the Oklahoma Medical Research Foundation and Wormbase. We further thank Lena Schumacher for excellent technical assistance. RNA sequencing was performed at the Genomics & Transcriptomics Laboratory (GTL) of the Center for Biological-Medical Research (BMFZ) at Heinrich Heine University Düsseldorf (BMFZ). Computational infrastructure and support were provided by the Centre for Information and Media Technology (ZIM) at Heinrich-Heine-University Düsseldorf. We thank Lena Abbey (Düsseldorf, Germany) for excellent technical assistance and Björn Schumacher (Cologne, Germany) for very helpful suggestions and discussion.

## Author contributions

JK: Conzeptualization; Investigation; Formal analysis; Methodology; Visualization; Writing-original draft

EI: Investigation

AP: Formal analysis

AKW: Investigation; Formal analysis; Visualization

TW: Investigation; Formal analysis

JB: Investigation; Formal analysis

SP: Supervision, Writing-original draft

GF: Conceptualization; Funding; Resources; Supervision; Writing-original draft; Writing-revision

## Abbreviations*

CI, chemotaxis Index; CIPN, chemotherapy-induced peripheral neuropathy; CisPt, cisplatin; DAVID, Database for Annotation, Visualization and Integrated Discovery; DDR, DNA damage response; DEG, differentially expressed gene; DRG, dorsal root ganglion; FDR, false discovery rate; GABA, gamma-aminobutyric acid; GFP, green fluorescent protein; GG-NER, global genome nucleotide excision repair; GO, Gene Onthology; GPCR, G protein-coupled receptor; KEGG, Kyoto Encyclopedia of Genes and Genomes; NER, nucleotide excision repair; NGM, nematode growth medium; RNA-seq, RNA sequencing; TC-NER, transcription-coupled nucleotide excision repair; UV-C, ultraviolet C light.

## Footnotes

* *C. elegans* genes are not included in this list. Their official WormBase names and identifiers are provided in Supplementary Tables S1 and S2.

## References

1. Cohen SM, Lippard SJ. Cisplatin: from DNA damage to cancer chemotherapy. Progress in nucleic acid research and molecular biology. 2001;67:93–130.

2. Reed E. Platinum-DNA adduct, nucleotide excision repair and platinum based anti-cancer chemotherapy. Cancer Treat Rev. 1998;24(5):331–344.

3. Hoeijmakers JH. Genome maintenance mechanisms for preventing cancer. Nature. 2001;411(6835):366–374.

4. Christmann M, Tomicic MT, Roos WP, Kaina B. Mechanisms of human DNA repair: an update. Toxicology. 2003;193(1-2):3–34.

5. Marteijn JA, Lans H, Vermeulen W, Hoeijmakers JH. Understanding nucleotide excision repair and its roles in cancer and ageing. Nat Rev Mol Cell Biol. 2014;15(7):465–481.

6. Wilson DM, Rieckher M, Williams AB, Schumacher B. Systematic analysis of DNA crosslink repair pathways during development and aging in Caenorhabditis elegans. Nucleic Acids Research. 2017;45(16):9467–9480.

7. van Sluis M, Gonzalo-Hansen C, Li Q, Lans H, Wang D, Marteijn JA. Mechanisms of transcription-coupled repair and DNA damage surveillance in health and disease. Nat Rev Mol Cell Biol. 2026;27(3):234–251.

8. Kumar V, Kashif M, Singh V, Rawat RS, Khan A, Sarkar GC, Majood M. Therapeutic targeting of DNA repair pathway dysregulation in aging, cancer, and neurodegeneration. Expert Opin Ther Targets. 2026:1–24.

9. Schumacher B, Pothof J, Vijg J, Hoeijmakers JHJ. The central role of DNA damage in the ageing process. Nature. 2021;592(7856):695–703.

10. Yao X, Panichpisal K, Kurtzman N, Nugent K. Cisplatin nephrotoxicity: a review. Am J Med Sci. 2007;334(2):115–124.

11. Patane S. Cardiotoxicity: cisplatin and long-term cancer survivors. Int J Cardiol. 2014;175(1):201–202.

12. Staff NP, Grisold A, Grisold W, Windebank AJ. Chemotherapy-Induced Peripheral Neuropathy: A Current Review. Ann Neurol. 2017;81(6):772–781.

13. Ezzi MS, Othieno-Abinya NA, Amayo E, Oyiro P, McLigeyo A, Yatich RB, Shoba B. Prevalence and Predictors of Cisplatin-Induced Peripheral Neuropathy at the Kenyatta National Hospital. J Glob Oncol. 2019;5:1–6.

14. Flatters SJL, Dougherty PM, Colvin LA. Clinical and preclinical perspectives on Chemotherapy-Induced Peripheral Neuropathy (CIPN): a narrative review. Br J Anaesth. 2017;119(4):737–749.

15. Salat K. Chemotherapy-induced peripheral neuropathy: part 1-current state of knowledge and perspectives for pharmacotherapy. Pharmacological Reports. 2020;72(3):486–507.

16. McDonald ES, Randon KR, Knight A, Windebank AJ. Cisplatin preferentially binds to DNA in dorsal root ganglion neurons in vitro and in vivo: a potential mechanism for neurotoxicity. Neurobiol Dis. 2005;18(2):305–313.

17. Estivill-Torrus G, Martinez-Padilla AB, Sanchez-Salido L, Evercooren AB, Garcia-Diaz B. The dorsal root ganglion as a target for neurorestoration in neuropathic pain. Neural Regen Res. 2024;19(2):296–301.

18. Huang Y, Chen H, Chen SR, Pan HL. Tonic GABA and Glycine Inhibition Control Pain Hypersensitivity via Limiting alpha2delta-1- and mGluR5-Dependent NMDA Receptor Activity at Primary Afferent-->Excitatory Neuron Synapses. J Neurosci. 2025;45(44).

19. Comitato A, Bardoni R. Presynaptic Inhibition of Pain and Touch in the Spinal Cord: From Receptors to Circuits. Int J Mol Sci. 2021;22(1).

20. Zeilhofer HU, Wildner H, Yevenes GE. Fast synaptic inhibition in spinal sensory processing and pain control. Physiol Rev. 2012;92(1):193–235.

21. Peirs C, Seal RP. Neural circuits for pain: Recent advances and current views. Science. 2016;354(6312):578–584.

22. Torsney C, MacDermott AB. Disinhibition opens the gate to pathological pain signaling in superficial neurokinin 1 receptor-expressing neurons in rat spinal cord. J Neurosci. 2006;26(6):1833–1843.

23. Cavaletti G, Marmiroli P. Chemotherapy-induced peripheral neurotoxicity. Nat Rev Neurol. 2010;6(12):657–666.

24. Velasco R, Bruna J. [Chemotherapy-induced peripheral neuropathy: an unresolved issue]. Neurologia. 2010;25(2):116–131.

25. Babbar M, Basu S, Yang B, Croteau DL, Bohr VA. Mitophagy and DNA damage signaling in human aging. Mech Ageing Dev. 2020;186:111207.

26. Madabhushi R, Pan L, Tsai LH. DNA damage and its links to neurodegeneration. Neuron. 2014;83(2):266–282.

27. Ahmad A, Braden A, Khan S, Xiao J, Khan MM. Crosstalk between the DNA damage response and cellular senescence drives aging and age-related diseases. Semin Immunopathol. 2024;46(3-4):10.

28. Rolig RL, McKinnon PJ. Linking DNA damage and neurodegeneration. Trends Neurosci. 2000;23(9):417–424.

29. Ropert B, Gallrein C, Schumacher B. DNA repair deficiencies and neurodegeneration. DNA Repair (Amst). 2024;138:103679.

30. Englander EW. DNA damage response in peripheral nervous system: coping with cancer therapy-induced DNA lesions. DNA Repair (Amst). 2013;12(8):685–690.

31. Acklin S, Xia F. The Role of Nucleotide Excision Repair in Cisplatin-Induced Peripheral Neuropathy: Mechanism, Prevention, and Treatment. Int J Mol Sci. 2021;22(4).

32. Honnen S. Caenorhabditis elegans as a powerful alternative model organism to promote research in genetic toxicology and biomedicine. Archives of Toxicology. 2017;91(5):2029–2044.

33. Leung MC, Williams PL, Benedetto A, Au C, Helmcke KJ, Aschner M, Meyer JN. Caenorhabditis elegans: an emerging model in biomedical and environmental toxicology. Toxicol Sci. 2008;106(1):5–28.

34. Garcia-Rodriguez FJ, Martinez-Fernandez C, Brena D, Kukhtar D, Serrat X, Nadal E, Boxem M, Honnen S, Miranda-Vizuete A, Villanueva A, Ceron J. Genetic and cellular sensitivity of Caenorhabditis elegans to the chemotherapeutic agent cisplatin. Disease Models & Mechanisms. 2018;11(6).

35. Rieckher M, Bujarrabal A, Doll MA, Soltanmohammadi N, Schumacher B. A simple answer to complex questions: Caenorhabditis elegans as an experimental model for examining the DNA damage response and disease genes. J Cell Physiol. 2018;233(4):2781–2790.

36. Lopes AFC, Bozek K, Herholz M, Trifunovic A, Rieckher M, Schumacher B. A C. elegans model for neurodegeneration in Cockayne syndrome. Nucleic Acids Research. 2020;48(19):10973–10985.

37. Wellenberg A, Brinkmann V, Bornhorst J, Ventura N, Honnen S, Fritz G. Cisplatin-induced neurotoxicity involves the disruption of serotonergic neurotransmission. Pharmacological Research. 2021;174.

38. Wellenberg A, Weides L, Kurzke J, Hennecke T, Bornhorst J, Crone B, Karst U, Brinkmann V, Fritz G, Honnen S. Use of C. elegans as a 3R-compliant in vivo model for the chemoprevention of cisplatin-induced neurotoxicity. Exp Neurol. 2021;341.

39. Martinez-Fernandez C, Bergamino M, Schiavi A, Brena D, Ventura N, Honnen S, Villanueva A, Nadal E, Ceron J. Insights into cisplatin-induced neurotoxicity and mitochondrial dysfunction in Caenorhabditis elegans. Dis Model Mech. 2022.

40. Lans H, Vermeulen W. Tissue specific response to DNA damage: C. elegans as role model. DNA Repair (Amst). 2015;32:141–148.

41. Lans H, Marteijn JA, Schumacher B, Hoeijmakers JH, Jansen G, Vermeulen W. Involvement of global genome repair, transcription coupled repair, and chromatin remodeling in UV DNA damage response changes during development. PLoS Genet. 2010;6(5):e1000941.

42. Sabatella M, Thijssen KL, Davo-Martinez C, Vermeulen W, Lans H. Tissue-Specific DNA Repair Activity of ERCC-1/XPF-1. Cell Reports. 2021;34(2).

43. Wynen F, Krautstrunk J, Muller LM, Graf V, Brinkmann V, Fritz G. Cisplatin-induced DNA crosslinks trigger neurotoxicity in C. elegans. Biochim Biophys Acta Mol Cell Res. 2024;1871(1):119591.

44. Consortium CeDM. large-scale screening for targeted knockouts in the Caenorhabditis elegans genome. G3 (Bethesda). 2012;2(11):1415–1425.

45. Stiernagle T. Maintenance of C. elegans. WormBook. 2006:1–11.

46. Xiao R, Chun L, Ronan EA, Friedman DI, Liu J, Xu XZ. RNAi Interrogation of Dietary Modulation of Development, Metabolism, Behavior, and Aging in C. elegans. Cell Rep. 2015;11(7):1123–1133.

47. Margie O, Palmer C, Chin-Sang I. C. elegans Chemotaxis Assay. Jove-J Vis Exp. 2013(74).

48. Chew YL, Fan X, Gotz J, Nicholas HR. Aging in the nervous system of Caenorhabditis elegans. Commun Integr Biol. 2013;6(5):e25288.

49. Love MI, Huber W, Anders S. Moderated estimation of fold change and dispersion for RNA-seq data with DESeq2. Genome Biol. 2014;15(12):550.

50. Stephens M. False discovery rates: a new deal. Biostatistics. 2017;18(2):275–294.

51. Huang da W, Sherman BT, Lempicki RA. Systematic and integrative analysis of large gene lists using DAVID bioinformatics resources. Nat Protoc. 2009;4(1):44–57.

52. Sherman BT, Panzade G, Dotrang T, Hao M, Xu L, Li X, Baseler MW, Lane HC, Imamichi T, Chang W. DAVID: a web server for functional annotation and functional enrichment analysis of gene lists (2025 update). Nucleic Acids Res. 2026.

53. Mahoney TR, Luo S, Nonet ML. Analysis of synaptic transmission in Caenorhabditis elegans using an aldicarb-sensitivity assay. Nat Protoc. 2006;1(4):1772–1777.

54. Weishaupt AK, Kubens L, Ruecker L, Schwerdtle T, Aschner M, Bornhorst J. A Reliable Method Based on Liquid Chromatography-Tandem Mass Spectrometry for the Simultaneous Quantification of Neurotransmitters in Caenorhabditis elegans. Molecules. 2023;28(14).

55. Bonhof CS, van de Poll-franse LV, Wasowicz DK, Beerepoot LV, Vreugdenhil G, Mols F. The course of peripheral neuropathy and its association with health-related quality of life among colorectal cancer patients. J Cancer Surviv. 2021;15(2):190–200.

56. Mounier NM, Abdel-Maged AES, Wahdan SA, Gad AM, Azab SS. Chemotherapy-induced cognitive impairment (CICI): An overview of etiology and pathogenesis. Life sciences. 2020;258.

57. Conte E, Bresciani E, Rizzi L, Cappellari O, De Luca A, Torsello A, Liantonio A. Cisplatin-Induced Skeletal Muscle Dysfunction: Mechanisms and Counteracting Therapeutic Strategies. Int J Mol Sci. 2020;21(4).

58. Maglioni S, Schiavi A, Runci A, Shaik A, Ventura N. Mitochondrial stress extends lifespan in C. elegans through neuronal hormesis. Experimental Gerontology. 2014;56:89–98.

59. Gurkar AU, Robinson AR, Cui Y, Li X, Allani SK, Webster A, Muravia M, Fallahi M, Weissbach H, Robbins PD, Wang Y, Kelley EE, Croix CMS, Niedernhofer LJ, Gill MS. Dysregulation of DAF-16/FOXO3A-mediated stress responses accelerates oxidative DNA damage induced aging. Redox Biol. 2018;18:191–199.

60. Niedernhofer LJ, Garinis GA, Raams A, Lalai AS, Robinson AR, Appeldoorn E, Odijk H, Oostendorp R, Ahmad A, van Leeuwen W, Theil AF, Vermeulen W, van der Horst GT, Meinecke P, Kleijer WJ, Vijg J, Jaspers NG, Hoeijmakers JH. A new progeroid syndrome reveals that genotoxic stress suppresses the somatotroph axis. Nature. 2006;444(7122):1038–1043.

61. Weeda G, Donker I, de Wit J, Morreau H, Janssens R, Vissers CJ, Nigg A, van Steeg H, Bootsma D, Hoeijmakers JHJ. Disruption of mouse ERCC1 results in a novel repair syndrome with growth failure, nuclear abnormalities and senescence. Curr Biol. 1997;7(6):427–439.

62. Vougioukalaki M, Demmers J, Vermeij WP, Baar M, Bruens S, Magaraki A, Kuijk E, Jager M, Merzouk S, Brandt RMC, Kouwenberg J, van Boxtel R, Cuppen E, Pothof J, Hoeijmakers JHJ. Different responses to DNA damage determine ageing differences between organs. Aging Cell. 2022;21(4):e13562.

63. West H, Coffey M, Wagner MJ, McLeod HL, Colley JP, Adams RA, Fleck O, Maughan TS, Fisher D, Kaplan RS, Harris R, Cheadle JP. Role for Nucleotide Excision Repair Gene Variants in Oxaliplatin-Induced Peripheral Neuropathy. JCO Precis Oncol. 2018;2:1–18.

64. Khrunin AV, Moisseev A, Gorbunova V, Limborska S. Genetic polymorphisms and the efficacy and toxicity of cisplatin-based chemotherapy in ovarian cancer patients. Pharmacogenomics J. 2010;10(1):54–61.

65. Quan Y, Zhang E, Nan Y, Lin H. Regional-specific GABAergic imbalance in cisplatin-induced neuropathy and antiallodynic effects of intraplantar baclofen in rats. Scientific reports. 2026;16(1).

66. da Silva PFL, Ropert B, Rieckher M, Rive C, Schumacher B. Neuronal DNA repair mediates systemic mitochondrial maintenance through acetylcholine signaling. bioRxiv. 2026:2026.2002.2011.705232.

67. Ijomone OM, Miah MR, Akingbade GT, Bucinca H, Aschner M. Nickel-Induced Developmental Neurotoxicity in C. elegans Includes Cholinergic, Dopaminergic and GABAergic Degeneration, Altered Behaviour, and Increased SKN-1 Activity. Neurotox Res. 2020;37(4):1018–1028.

68. Cavaletti G, Tredici G. Cisplatin neuropathy. Neurology. 1995;45(3 Pt 1):596–597.

69. Cheng H, Chen D, Li X, Al-Sheikh U, Duan D, Fan Y, Zhu L, Zeng W, Hu Z, Tong X, Zhao G, Zhang Y, Zou W, Duan S, Kang L. Phasic/tonic glial GABA differentially transduce for olfactory adaptation and neuronal aging. Neuron. 2024;112(9):1473–1486 e1476.

70. Wirak GS, Florman J, Alkema MJ, Connor CW, Gabel CV. Age-associated changes to neuronal dynamics involve a disruption of excitatory/inhibitory balance in C. elegans. Elife. 2022;11.

71. Wu J, Wang L, Ervin JF, Wang SJ, Soderblom E, Ko D, Yan D. GABA signaling triggered by TMC-1/Tmc delays neuronal aging by inhibiting the PKC pathway in C. elegans. Sci Adv. 2022;8(51):eadc9236.

72. McMillen A, Minervini C, Green R, Johnson ME, Ansaar R, Chew YL. Dopaminergic Modulation of Short-Term Associative Memory in Caenorhabditis elegans. J Neurochem. 2025;169(8):e70200.

73. Lans H, Jansen G. Multiple sensory G proteins in the olfactory, gustatory and nociceptive neurons modulate longevity in Caenorhabditis elegans. Dev Biol. 2007;303(2):474–482.

74. Murphy CT, McCarroll SA, Bargmann CI, Fraser A, Kamath RS, Ahringer J, Li H, Kenyon C. Genes that act downstream of DAF-16 to influence the lifespan of Caenorhabditis elegans. Nature. 2003;424(6946):277–283.

75. Ogg S, Paradis S, Gottlieb S, Patterson GI, Lee L, Tissenbaum HA, Ruvkun G. The Fork head transcription factor DAF-16 transduces insulin-like metabolic and longevity signals in C. elegans. Nature. 1997;389(6654):994–999.

76. Becker G, Atuati SF, Oliveira SM. G Protein-Coupled Receptors and Ion Channels Involvement in Cisplatin-Induced Peripheral Neuropathy: A Review of Preclinical Studies. Cancers (Basel). 2024;16(3).

77. Uniyal A, Tiwari V, Tsukamoto T, Dong X, Guan Y, Raja SN. Targeting sensory neuron GPCRs for peripheral neuropathic pain. Trends Pharmacol Sci. 2023;44(12):1009–1027.

