## Supplementary Tables for "Cisplatin-triggered sensory dysfunction in NER-defective *C. elegans* involves GABAergic and cholinergic signaling"

**Supplementary Table S1: Genes examined or referenced throughout the study.**

Official WormBase names and identifiers of genes investigated using mutant strains or presented individually in Fig. 1E.

| **Gene** | **Full name (WormBase)** | **WormBase ID** |
| --- | --- | --- |
| *ace-1* | abnormal AcetylCholinEsterase 1 | WBGene00000035 |
| *ace-2* | abnormal AcetylCholinEsterase 2 | WBGene00000036 |
| *acr-2* | AcetylCholine Receptor 2 | WBGene00000042 |
| *atl-1* | ATM (ataxia telangectasia mutated)-Like 1 | WBGene00000226 |
| *atm-1* | ATM (ataxia telangectasia mutated) family 1 | WBGene00000227 |
| *cep-1* | C. Elegans P-53-like protein 1 | WBGene00000467 |
| *cha-1* | abnormal CHoline Acetyltransferase 1 | WBGene00000481 |
| *cho-1* | CHOline transporter 1 | WBGene00000501 |
| *clk-1* | CLocK (biological timing) abnormality 1 | WBGene00000536 |
| *daf-16* | abnormal DAuer Formation 16 | WBGene00000912 |
| *daf-2* | abnormal DAuer Formation 2 | WBGene00000898 |
| *dgk-1* | DiacylGlycerol Kinase 1 | WBGene00000958 |
| *dlk-1* | Dual-Leucine zipper Kinase 1 | WBGene00001008 |
| *eat-2* | EATing: abnormal pharyngeal pumping 2 | WBGene00001133 |
| *ercc-1* | ERCC (DNA excision repair protein) homolog 1 | WBGene00008665 |
| *exp-1* | EXPulsion defective (defecation) 1 | WBGene00001373 |
| *fbl-1* | FiBuLin 1 | WBGene00001403 |
| *gar-3* | G-protein-linked Acetylcholine Receptor 3 | WBGene00001519 |
| *gbb-1* | GABA B receptor subunit 1 | WBGene00021528 |
| *gbb-2* | GABA B receptor subunit 2 | WBGene00022675 |
| *him-4* | High Incidence of Males (increased X chromosome loss) 4 | WBGene00001863 |
| *hsf-1* | Heat Shock Factor 1 | WBGene00002004 |
| *jkk-1* | JNK Kinase 1 | WBGene00002177 |
| *jnk-1* | Jun N-terminal Kinase 1 | WBGene00002178 |
| *lgc-36* | Ligand-Gated ion Channel 36 | WBGene00008551 |
| *lgc-37* | Ligand-Gated ion Channel 37 | WBGene00013914 |
| *lin-15* | abnormal cell LINeage 15 | WBGene00003004 |
| *lmn-1* | nuclear LaMiN 1 | WBGene00003052 |
| *mec-1* | MEChanosensory abnormality 1 | WBGene00003165 |
| *mec-10* | MEChanosensory abnormality 10 | WBGene00003174 |
| *mec-12* | MEChanosensory abnormality 12 | WBGene00003175 |
| *mec-2* | MEChanosensory abnormality 2 | WBGene00003166 |
| *mec-4* | MEChanosensory abnormality 4 | WBGene00003168 |
| *mec-5* | MEChanosensory abnormality 5 | WBGene00003169 |
| *mec-6* | MEChanosensory abnormality 6 | WBGene00003170 |
| *mec-9* | MEChanosensory abnormality 9 | WBGene00003173 |
| *mek-1* | MAP kinase kinase or Erk Kinase 1 | WBGene00003185 |
| *mev-1* | abnormal MEthyl Viologen sensitivity 1 | WBGene00003225 |
| *mkk-4* | MKK (MAP Kinase Kinase) homolog 4 | WBGene00003368 |
| *mlk-1* | human MLK (Mixed Lineage Kinase) homolog 1 | WBGene00003374 |
| *nsy-1* | Neuronal SYmmetry 1 | WBGene00003822 |
| *odr-10* | ODoRant response abnormal 10 | WBGene00003856 |
| *pmk-3* | P38 Map Kinase family 3 | WBGene00004057 |
| *ptl-1* | Protein with Tau-Like repeats 1 | WBGene00004212 |
| *sek-1* | SAPK/ERK kinase 1 | WBGene00004758 |
| *slo-1* | SLOwpoke potassium channel family 1 | WBGene00004830 |
| *unc-104* | UNCoordinated 104 | WBGene00006831 |
| *unc-25* | UNCoordinated 25 | WBGene00006762 |
| *unc-47* | UNCoordinated 47 | WBGene00006783 |
| *unc-49* | UNCoordinated 49 | WBGene00006784 |

**Supplementary Table S2: Genes represented in the RNA-seq functional annotation clusters.**

Official WormBase names and identifiers of genes represented in the functional annotation clusters shown in Supplementary Figs. S6-S8.

| **Gene** | **Full name (WormBase)** | **WormBase ID** |
| --- | --- | --- |
| *abt-1* | ABC Transporter family 1 | WBGene00000019 |
| *abt-5* | ABC Transporter family 5 | WBGene00000023 |
| *acc-3* | Acetylcholine-gated Chloride Channel 3 | WBGene00018880 |
| *acr-12* | AcetylCholine Receptor 12 | WBGene00000051 |
| *acr-19* | AcetylCholine Receptor 19 | WBGene00000058 |
| *acr-2* | AcetylCholine Receptor 2 | WBGene00000042 |
| *acr-25* | AcetylCholine Receptor 25 | WBGene00043066 |
| *acs-18* | fatty Acid CoA Synthetase family 18 | WBGene00011173 |
| *acy-1* | Adenylyl CYclase 1 | WBGene00000068 |
| *acy-2* | Adenylyl CYclase 2 | WBGene00000069 |
| *ador-1* | ADenOsine Receptor homolog 1 | WBGene00011878 |
| *aex-3* | ABoc, EXpulsion (defecation) defective 3 | WBGene00000086 |
| *aptf-1* | AP-2 Transcription Factor family 1 | WBGene00019424 |
| *ast-1* | Axon STeering defect 1 | WBGene00020368 |
| *avr-14* | altered AVeRmectin sensitivity 14 | WBGene00000232 |
| *B0207.7* | Protein kinase domain-containing protein | WBGene00015030 |
| *B0218.5* | Protein kinase domain-containing protein | WBGene00015049 |
| *bbs-2* | BBS (Bardet-Biedl Syndrome) protein 2 | WBGene00000242 |
| *bbs-8* | BBS (Bardet-Biedl Syndrome) protein 8 | WBGene00000244 |
| *bcl-11* | mammalian BCL (B cell lymphoma) gene homologs 11 | WBGene00017430 |
| *bed-3* | BED-type zinc finger putative transcription factor 3 | WBGene00009133 |
| *best-23* | BESTrophin (chloride channel) homolog 23 | WBGene00014989 |
| *blmp-1* | Blimp1 (B Lymphocyte-induced Maturation Protein-1) homolog 1 | WBGene00003847 |
| *bro-1* | BROther (Drosophila tx factor partner) homolog 1 | WBGene00000272 |
| *C05C12.1* | Protein kinase domain-containing protein | WBGene00007335 |
| *C08F8.6* | non-specific serine/threonine protein kinase | WBGene00007448 |
| *C09B9.4* | defective SPErmatogenesis 55 | WBGene00015629 |
| *C09D4.3* | Protein kinase domain-containing protein | WBGene00015634 |
| *C10G11.8* | AAA+ ATPase domain-containing protein | WBGene00015688 |
| *C16E9.1* | VWFA domain-containing protein | WBGene00015865 |
| *C17C3.11* | Protein kinase domain-containing protein | WBGene00015893 |
| *C18H7.4* | Tyrosine-protein kinase | WBGene00015994 |
| *C25A8.5* | Tyrosine-protein kinase | WBGene00016085 |
| *C26B2.2* | Membrane-associated kinase regulator | WBGene00016125 |
| *C27D6.11* | Protein kinase domain-containing protein | WBGene00044388 |
| *C27D8.1* | Protein kinase domain-containing protein | WBGene00007777 |
| *C28A5.6* | Protein kinase domain-containing protein | WBGene00007791 |
| *C33F10.8* | Tyrosine-protein phosphatase domain-containing protein | WBGene00016356 |
| *C34B2.3* | Protein kinase domain-containing protein | WBGene00016388 |
| *C34B4.3* | G_PROTEIN_RECEP_F1_2 domain-containing protein | WBGene00007908 |
| *C34E11.2* | MH2 domain-containing protein | WBGene00007935 |
| *C34F11.5* | Tyrosine-protein kinase | WBGene00016416 |
| *C35E7.10* | Tyrosine-protein kinase | WBGene00016462 |
| *C36E8.4* | SH3 domain-containing protein | WBGene00007983 |
| *C39B10.1* | G-protein coupled receptors family 1 profile domain-containing protein | WBGene00008021 |
| *C49C8.1* | Protein kinase domain-containing protein | WBGene00016765 |
| *C50H2.4* | Homeobox domain-containing protein | WBGene00008240 |
| *C50H2.7* | Protein kinase domain-containing protein | WBGene00008243 |
| *C52E4.7* | Histidine kinase | WBGene00008259 |
| *C55B7.10* | Protein kinase domain-containing protein | WBGene00016946 |
| *C55C3.4* | Tyrosine-protein kinase | WBGene00016954 |
| *C56C10.6* | Protein kinase domain-containing protein | WBGene00016963 |
| *catp-3* | Cation transporting ATPase 3 | WBGene00015660 |
| *catp-4* | Cation transporting ATPase 4 | WBGene00007248 |
| *ceh-17* | C. Elegans Homeobox 17 | WBGene00000440 |
| *ceh-2* | C. Elegans Homeobox 2 | WBGene00000429 |
| *ceh-20* | C. Elegans Homeobox 20 | WBGene00000443 |
| *ceh-22* | C. Elegans Homeobox 22 | WBGene00000445 |
| *ceh-30* | C. Elegans Homeobox 30 | WBGene00000451 |
| *ceh-31* | C. Elegans Homeobox 31 | WBGene00000452 |
| *ceh-32* | C. Elegans Homeobox 32 | WBGene00000453 |
| *ceh-33* | C. Elegans Homeobox 33 | WBGene00000454 |
| *ceh-34* | C. Elegans Homeobox 34 | WBGene00000455 |
| *ceh-45* | C. Elegans Homeobox 45 | WBGene00022837 |
| *ceh-48* | C. Elegans Homeobox 48 | WBGene00015934 |
| *ceh-6* | C. Elegans Homeobox 6 | WBGene00000431 |
| *ces-2* | CEll death Specification 2 | WBGene00000469 |
| *cest-16* | Carboxyl ESTerase domain containing 16 | WBGene00020330 |
| *cest-17* | Carboxyl ESTerase domain containing 17 | WBGene00019477 |
| *cest-19* | Carboxyl ESTerase domain containing 19 | WBGene00020125 |
| *cfi-1* | CEM Fate Inhibitor 1 | WBGene00000476 |
| *cky-1* | CKY homolog 1 | WBGene00000521 |
| *cla-1* | CLArinet, functional homolog of cytomatrix at the active zone proteins piccolo and fife 1 | WBGene00018468 |
| *clhm-1* | CaLcium Homeostasis Modulator 1 | WBGene00016626 |
| *cmk-1* | CaM Kinase 1 | WBGene00000553 |
| *cng-2* | Cyclic Nucleotide Gated channel 2 | WBGene00022295 |
| *cog-1* | Connection Of Gonad defective 1 | WBGene00000584 |
| *comp-1* | sperm COMPetition 1 | WBGene00018158 |
| *ctbp-1* | CTBP (CtBP) transcriptional co-repressor homolog 1 | WBGene00006424 |
| *D2045.5* | non-specific serine/threonine protein kinase | WBGene00008423 |
| *D2062.4* | Histidine kinase | WBGene00017056 |
| *dat-1* | DopAmine Transporter 1 | WBGene00000934 |
| *degt-1* | DEGenerin-related Touch-involved channel 1 | WBGene00009109 |
| *del-7* | DEgenerin Like 7 | WBGene00016699 |
| *dgk-1* | DiacylGlycerol Kinase 1 | WBGene00000958 |
| *dlg-1* | Drosophila Discs LarGe homolog 1 | WBGene00001006 |
| *dmd-10* | DM (Doublesex/MAB-3) Domain family 10 | WBGene00007929 |
| *dmsr-11* | DroMyoSuppressin Receptor related 11 | WBGene00022605 |
| *dmsr-5* | DroMyoSuppressin Receptor related 5 | WBGene00012962 |
| *dmsr-7* | DroMyoSuppressin Receptor related 7 | WBGene00016428 |
| *dop-5* | DOPamine receptor 5 | WBGene00011382 |
| *dop-6* | DOPamine receptor 6 | WBGene00016037 |
| *dyb-1* | DYstroBrevin homolog 1 | WBGene00001115 |
| *eat-16* | EATing: abnormal pharyngeal pumping 16 | WBGene00001145 |
| *eat-18* | EATing: abnormal pharyngeal pumping 18 | WBGene00001147 |
| *eat-2* | EATing: abnormal pharyngeal pumping 2 | WBGene00001133 |
| *egl-2* | EGg Laying defective 2 | WBGene00001171 |
| *egl-38* | EGg Laying defective 38 | WBGene00001204 |
| *egl-43* | EGg Laying defective 43 | WBGene00001207 |
| *elt-4* | Erythroid-Like Transcription factor family 4 | WBGene00001252 |
| *exp-2* | EXPulsion defective (defecation) 2 | WBGene00001374 |
| *eyg-1* | EYG (Drosophila eyegone) homolog 1 | WBGene00013147 |
| *F10G8.2* | Protein kinase domain-containing protein | WBGene00008662 |
| *F20H11.4* | Protein-tyrosine phosphatase | WBGene00017647 |
| *F21F3.2* | Protein kinase domain-containing protein | WBGene00017672 |
| *F22B3.8* | Tyrosine-protein kinase | WBGene00009039 |
| *F25F2.1* | Protein kinase domain-containing protein | WBGene00009115 |
| *F26A1.3* | non-specific serine/threonine protein kinase | WBGene00017802 |
| *F26A1.4* | non-specific serine/threonine protein kinase | WBGene00017803 |
| *F26A10.2* | C2H2-type domain-containing protein | WBGene00017814 |
| *F26E4.5* | Tyrosine-protein kinase | WBGene00009160 |
| *F32B5.1* | arginine kinase | WBGene00017975 |
| *F33D11.7* | Protein kinase domain-containing protein | WBGene00018004 |
| *F36H12.9* | Protein kinase domain-containing protein | WBGene00018123 |
| *F41G3.5* | non-specific serine/threonine protein kinase | WBGene00018301 |
| *F47E1.20* | C2H2-type domain-containing protein | WBGene00304808 |
| *F53C3.1* | Protein kinase domain-containing protein | WBGene00018745 |
| *F54H5.2* | Protein kinase domain-containing protein | WBGene00018839 |
| *F59A3.8* | Tyrosine-protein kinase | WBGene00019081 |
| *F59A6.4* | Protein kinase domain-containing protein | WBGene00019086 |
| *F59E12.3* | Protein kinase domain-containing protein | WBGene00019119 |
| *F59F3.6* | Potassium channel tetramerisation-type BTB domain-containing protein | WBGene00010335 |
| *feh-1* | mammalian FE65 Homolog 1 | WBGene00001410 |
| *fipr-24* | FIP (Fungus-Induced Protein) Related 24 | WBGene00007992 |
| *frk-1* | Fer oncogene Related Kinase 1 | WBGene00001487 |
| *frm-2* | FERM domain (protein4.1-ezrin-radixin-moesin) family 2 | WBGene00001489 |
| *gab-1* | GABa receptor subunit 1 | WBGene00001512 |
| *gar-2* | G-protein-linked Acetylcholine Receptor 2 | WBGene00001518 |
| *gar-3* | G-protein-linked Acetylcholine Receptor 3 | WBGene00001519 |
| *gbb-2* | GABA B receptor subunit 2 | WBGene00022675 |
| *gcy-29* | Guanylyl CYclase 29 | WBGene00007314 |
| *glc-3* | Glutamate-gated ChLoride channel 3 | WBGene00001593 |
| *glc-4* | Glutamate-gated ChLoride channel 4 | WBGene00001594 |
| *gldi-10* | Germ Layers DIsorganized 10 | WBGene00007222 |
| *gln-2* | GLutamiNe synthetase (glutamate-ammonia ligase) 2 | WBGene00001603 |
| *glr-1* | GLutamate Receptor family (AMPA) 1 | WBGene00001612 |
| *glr-5* | GLutamate Receptor family (AMPA) 5 | WBGene00001616 |
| *glr-7* | GLutamate Receptor family (AMPA) 7 | WBGene00001618 |
| *glt-5* | GLutamate Transporter family 5 | WBGene00001623 |
| *gpa-11* | G Protein, Alpha subunit 11 | WBGene00001673 |
| *gpa-2* | G Protein, Alpha subunit 2 | WBGene00001664 |
| *grh-1* | GRainyHead (Drosophila transcription factor) homolog 1 | WBGene00001707 |
| *gskl-1* | GSK3 Like 1 | WBGene00010991 |
| *gskl-2* | GSK3 Like 2 | WBGene00007977 |
| *H05L14.1* | Protein kinase domain-containing protein | WBGene00010366 |
| *hasp-2* | HASPin kinase related 2 | WBGene00021214 |
| *his-10* | HIStone 10 | WBGene00001884 |
| *hlh-10* | Helix Loop Helix 10 | WBGene00001954 |
| *hlh-13* | Helix Loop Helix 13 | WBGene00001957 |
| *hlh-32* | Helix Loop Helix 32 | WBGene00013665 |
| *hlh-34* | Helix Loop Helix 34 | WBGene00011327 |
| *hlh-6* | Helix Loop Helix 6 | WBGene00001952 |
| *hsp-16.2* | Heat Shock Protein 16.2 | WBGene00002016 |
| *hsp-16.20* | Heat Shock Protein 16.20 | WBGene00008591 |
| *hsp-16.41* | Heat Shock Protein 16.41 | WBGene00002018 |
| *hsp-16.48* | Heat Shock Protein 16.48 | WBGene00002019 |
| *hsp-16.49* | Heat Shock Protein 16.49 | WBGene00002020 |
| *hsp-70* | Heat Shock Protein 70 | WBGene00002026 |
| *ins-1* | INSulin related 1 | WBGene00002084 |
| *ins-20* | INSulin related 20 | WBGene00002103 |
| *ins-31* | INSulin related 31 | WBGene00002114 |
| *ins-37* | INSulin related 37 | WBGene00002120 |
| *inx-18* | INneXin 18 | WBGene00002140 |
| *jmjc-1* | JuManJiC domain protein 1 | WBGene00020902 |
| *jmjd-3.1* | JuMonJi (transcription factor) Domain protein 3.1 | WBGene00017571 |
| *K06H7.8* | Putative serine/threonine-protein kinase K06H7.1 | WBGene00019459 |
| *K07A3.3* | ATP-dependent Clp protease ATP-binding subunit clpX-like, mitochondrial | WBGene00019461 |
| *K09C6.2* | Protein kinase domain-containing protein | WBGene00019556 |
| *K09C6.7* | Protein kinase domain-containing protein | WBGene00019561 |
| *K09E4.1* | Protein kinase domain-containing protein | WBGene00010719 |
| *K11C4.1* | non-specific serine/threonine protein kinase | WBGene00019642 |
| *kin-21* | protein KINase 21 | WBGene00002204 |
| *kin-24* | protein KINase 24 | WBGene00002206 |
| *kin-26* | protein KINase 26 | WBGene00002208 |
| *kin-5* | protein KINase 5 | WBGene00002193 |
| *klf-2* | Kruppel-Like Factor (zinc finger protein) 2 | WBGene00009998 |
| *klp-17* | Kinesin-Like Protein 17 | WBGene00002227 |
| *kpc-1* | Kex-2 Proprotein Convertase family 1 | WBGene00002232 |
| *kvs-3* | K (potassium) Voltage-Sensitive channel subunit 3 | WBGene00018402 |
| *lad-2* | L1 CAM ADhesion molecule homolog 2 | WBGene00002243 |
| *lea-1* | plant Late Embryo Abundant (LEA) related 1 | WBGene00002263 |
| *let-381* | LEThal 381 | WBGene00002601 |
| *lev-1* | LEVamisole resistant 1 | WBGene00002974 |
| *lgc-11* | Ligand-Gated ion Channel 11 | WBGene00000061 |
| *lgc-13* | Ligand-Gated ion Channel 13 | WBGene00011354 |
| *lgc-21* | Ligand-Gated ion Channel 21 | WBGene00007903 |
| *lgc-23* | Ligand-Gated ion Channel 23 | WBGene00007594 |
| *lgc-27* | Ligand-Gated ion Channel 27 | WBGene00022283 |
| *lgc-31* | Ligand-Gated ion Channel 31 | WBGene00008997 |
| *lgc-33* | Ligand-Gated ion Channel 33 | WBGene00021941 |
| *lgc-39* | Ligand-Gated ion Channel 39 | WBGene00017314 |
| *lgc-43* | Ligand-Gated ion Channel 43 | WBGene00008076 |
| *lgc-47* | Ligand-Gated ion Channel 47 | WBGene00009799 |
| *lgc-49* | Ligand-Gated ion Channel 49 | WBGene00010741 |
| *lgc-51* | Ligand-Gated ion Channel 51 | WBGene00017399 |
| *lgc-55* | Ligand-Gated ion Channel 55 | WBGene00013746 |
| *lgc-57* | Ligand-Gated ion Channel 57 | WBGene00001586 |
| *lgc-6* | Ligand-Gated ion Channel 6 | WBGene00017544 |
| *lgc-8* | Ligand-Gated ion Channel 8 | WBGene00013338 |
| *lin-22* | abnormal cell LINeage 22 | WBGene00003008 |
| *lys-3* | LYSozyme 3 | WBGene00003092 |
| *M04G7.2* | Protein-tyrosine phosphatase | WBGene00019773 |
| *M176.9* | Tyrosine-protein kinase | WBGene00014813 |
| *M7.7* | Protein kinase domain-containing protein | WBGene00010883 |
| *madd-4* | Muscle Arm Development Defective 4 | WBGene00009958 |
| *mec-2* | MEChanosensory abnormality 2 | WBGene00003166 |
| *mec-4* | MEChanosensory abnormality 4 | WBGene00003168 |
| *mls-2* | Mesodermal Lineage Specification 2 | WBGene00003377 |
| *nca-2* | Novel Channel type/putative Nematode CAlcium channel 2 | WBGene00003558 |
| *ncx-5* | Na/Ca eXchangers 5 | WBGene00003570 |
| *nhr-117* | Nuclear Hormone Receptor family 117 | WBGene00003707 |
| *nhr-120* | Nuclear Hormone Receptor family 120 | WBGene00003710 |
| *nhr-124* | Nuclear Hormone Receptor family 124 | WBGene00003714 |
| *nhr-136* | Nuclear Hormone Receptor family 136 | WBGene00003726 |
| *nhr-140* | Nuclear Hormone Receptor family 140 | WBGene00016366 |
| *nhr-147* | Nuclear Hormone Receptor family 147 | WBGene00015395 |
| *nhr-172* | Nuclear Hormone Receptor family 172 | WBGene00016927 |
| *nhr-18* | Nuclear Hormone Receptor family 18 | WBGene00003617 |
| *nhr-201* | Nuclear Hormone Receptor family 201 | WBGene00019741 |
| *nhr-208* | Nuclear Hormone Receptor family 208 | WBGene00011099 |
| *nhr-221* | Nuclear Hormone Receptor family 221 | WBGene00020748 |
| *nhr-25* | Nuclear Hormone Receptor family 25 | WBGene00003623 |
| *nhr-262* | Nuclear Hormone Receptor family 262 | WBGene00008619 |
| *nhr-275* | Nuclear Hormone Receptor family 275 | WBGene00021163 |
| *nhr-37* | Nuclear Hormone Receptor family 37 | WBGene00018412 |
| *nhr-41* | Nuclear Hormone Receptor family 41 | WBGene00022423 |
| *nhr-52* | Nuclear Hormone Receptor family 52 | WBGene00003642 |
| *nhr-54* | Nuclear Hormone Receptor family 54 | WBGene00003644 |
| *nhr-58* | Nuclear Hormone Receptor family 58 | WBGene00003648 |
| *nhr-59* | Nuclear Hormone Receptor family 59 | WBGene00003649 |
| *nhr-63* | Nuclear Hormone Receptor family 63 | WBGene00003653 |
| *nhr-67* | Nuclear Hormone Receptor family 67 | WBGene00003657 |
| *nhr-91* | Nuclear Hormone Receptor family 91 | WBGene00003681 |
| *nhr-99* | Nuclear Hormone Receptor family 99 | WBGene00003689 |
| *nlg-1* | NeuroLiGin family 1 | WBGene00006412 |
| *nmr-2* | NMDA class glutamate Receptor 2 | WBGene00003775 |
| *npax-2* | N-terminal PAX (PAI domain only) protein 2 | WBGene00018591 |
| *npr-14* | NeuroPeptide Receptor family 14 | WBGene00012275 |
| *npr-2* | NeuroPeptide Receptor family 2 | WBGene00003808 |
| *npr-23* | NeuroPeptide Receptor family 23 | WBGene00021328 |
| *npr-26* | NeuroPeptide Receptor family 26 | WBGene00011372 |
| *npr-3* | NeuroPeptide Receptor family 3 | WBGene00007006 |
| *npr-32* | NeuroPeptide Receptor family 32 | WBGene00013782 |
| *npr-38* | NeuroPeptide Receptor family 38 | WBGene00020420 |
| *npr-40* | NeuroPeptide Receptor family 40 | WBGene00013496 |
| *npr-41* | NeuroPeptide Receptor family 41 | WBGene00007146 |
| *npr-7* | NeuroPeptide Receptor family 7 | WBGene00018067 |
| *nrx-1* | NeuReXin related 1 | WBGene00007799 |
| *odd-2* | Drosophila ODD-skipped-like 2 | WBGene00003846 |
| *odr-3* | ODoRant response abnormal 3 | WBGene00003850 |
| *osm-5* | OSMotic avoidance abnormal 5 | WBGene00003885 |
| *pag-3* | PAttern of reporter Gene expression abnormal 3 | WBGene00003909 |
| *pat-9* | Paralysed Arrest at Two-fold 9 | WBGene00003933 |
| *pcdr-1* | Pathogen Clearance Defective Receptor 1 | WBGene00010329 |
| *pde-1* | PhosphoDiEsterase 1 | WBGene00011433 |
| *peb-1* | Pharyngeal Enhancer Binding 1 | WBGene00003968 |
| *pfk-1.2* | 6-PhosphoFructoKinase (E.C. 2.7.1.11) homolog 1.2 | WBGene00008230 |
| *pha-2* | defective PHArynx development 2 | WBGene00004011 |
| *pha-4* | defective PHArynx development 4 | WBGene00004013 |
| *ptl-1* | Protein with Tau-Like repeats 1 | WBGene00004212 |
| *ptrg-1* | Protein Tyrosine phosphatase Receptor type G 1 | WBGene00010558 |
| *R03H10.7* | Replication protein A OB domain-containing protein | WBGene00019859 |
| *R10D12.10* | Protein kinase domain-containing protein | WBGene00011191 |
| *R11E3.1* | Tyrosine-protein kinase | WBGene00020003 |
| *R13H7.2* | G-protein coupled receptors family 1 profile domain-containing protein | WBGene00020069 |
| *rab-3* | RAB family 3 | WBGene00004267 |
| *rad-50* | RADiation sensitivity abnormal/yeast RAD-related 50 | WBGene00004296 |
| *rbf-1* | RaBPHilin 1 | WBGene00004316 |
| *ric-4* | Resistance to Inhibitors of Cholinesterase 4 | WBGene00004364 |
| *rig-5* | neuRonal IGCAM 5 | WBGene00004372 |
| *rimb-1* | RIM Binding protein 1 | WBGene00006513 |
| *rnt-1* | RuNT related 1 | WBGene00004393 |
| *rpm-1* | Regulator of Presynaptic Morphology 1 | WBGene00004457 |
| *rsbp-1* | R-Seven Binding Protein (R7BP) homolog 1 | WBGene00011531 |
| *rskn-2* | RSK-pNinety (RSK-p90 kinase) homolog 2 | WBGene00008311 |
| *sad-1* | Synapses of Amphids Defective 1 | WBGene00004719 |
| *sdf-9* | Synthetic Dauer Formation 9 | WBGene00004748 |
| *ser-2* | SERotonin/octopamine receptor family 2 | WBGene00004777 |
| *shl-1* | SHaL family of potassium channels 1 | WBGene00022240 |
| *shn-1* | SHaNk (SH3/ankyrin domain scaffold protein) related 1 | WBGene00006444 |
| *shw-3* | SHaW family of potassium channels 3 | WBGene00004793 |
| *snf-11* | Sodium: Neurotransmitter symporter Family 11 | WBGene00004910 |
| *snt-1* | SyNapTotagmin 1 | WBGene00004921 |
| *sol-1* | Suppressor Of Lurcher movement defect 1 | WBGene00004944 |
| *somi-1* | Suppressor of Overexpressed MIcro-RNA 1 | WBGene00010868 |
| *sorb-1* | SORBin homolog 1 | WBGene00012891 |
| *sox-2* | SOX (mammalian SRY box) family 2 | WBGene00004949 |
| *sox-3* | SOX (mammalian SRY box) family 3 | WBGene00004950 |
| *spe-8* | defective SPErmatogenesis 8 | WBGene00004962 |
| *sphr-1* | SPHingosine-1-phosphate Receptor functional homolog 1 | WBGene00016044 |
| *sprr-1* | Sex Peptide Receptor (Drosophila) Related 1 | WBGene00010986 |
| *srg-30* | Serpentine Receptor, class G (gamma) 30 | WBGene00005187 |
| *srw-12* | Serpentine Receptor, class W 12 | WBGene00005759 |
| *srw-71* | Serpentine Receptor, class W 71 | WBGene00005818 |
| *srw-85* | Serpentine Receptor, class W 85 | WBGene00005832 |
| *srw-87* | Serpentine Receptor, class W 87 | WBGene00005834 |
| *stn-2* | SynTrophiN 2 | WBGene00017866 |
| *str-112* | Seven TM Receptor 112 | WBGene00006163 |
| *sup-9* | SUPpressor 9 | WBGene00006318 |
| *syd-1* | SYnapse Defective 1 | WBGene00006363 |
| *syg-1* | SYnaptoGenesis abnormal 1 | WBGene00006365 |
| *T05A7.6* | Protein kinase domain-containing protein | WBGene00020223 |
| *T05C12.1* | Protein kinase domain-containing protein | WBGene00011466 |
| *T06C10.3* | Tyrosine-protein kinase | WBGene00020289 |
| *T11F8.4* | non-specific serine/threonine protein kinase | WBGene00020435 |
| *T11F9.1* | G-protein coupled receptors family 1 profile domain-containing protein | WBGene00011709 |
| *T15B12.2* | non-specific serine/threonine protein kinase | WBGene00020531 |
| *T19D12.5* | non-specific serine/threonine protein kinase | WBGene00020580 |
| *T21G5.1* | Tyrosine-protein kinase | WBGene00020659 |
| *T25B9.4* | Tyrosine-protein kinase | WBGene00012010 |
| *T25B9.6* | Protein kinase domain-containing protein | WBGene00012012 |
| *T25F10.4* | Neur_chan_LBD domain-containing protein | WBGene00020807 |
| *T27E4.1* | C2 domain-containing protein | WBGene00020861 |
| *T28F4.3* | Protein-tyrosine phosphatase | WBGene00012138 |
| *tag-180* | Temporarily Assigned Gene name 180 | WBGene00007041 |
| *tag-191* | Temporarily Assigned Gene name 191 | WBGene00007049 |
| *tag-344* | Temporarily Assigned Gene name 344 | WBGene00015230 |
| *tax-2* | abnormal CHEmotaxis 2 | WBGene00006525 |
| *tkr-2* | TachyKinin Receptor family 2 | WBGene00016761 |
| *tmc-1* | TransMembrane Channel-like protein family 1 | WBGene00020490 |
| *trp-1* | TRP (transient receptor potential) channel family 1 | WBGene00006614 |
| *trpl-4* | TRP-channel-Like 4 | WBGene00021408 |
| *ttbk-2.1* | Tau TuBulin Kinase 2.1 | WBGene00018122 |
| *ttbk-2.2* | Tau TuBulin Kinase 2.2 | WBGene00020071 |
| *ttbk-3* | Tau TuBulin Kinase 3 | WBGene00009324 |
| *ttbk-4* | Tau TuBulin Kinase 4 | WBGene00012169 |
| *ttbk-5* | Tau TuBulin Kinase 5 | WBGene00007305 |
| *ttbk-8.1* | Tau TuBulin Kinase 8.1 | WBGene00012637 |
| *ttbk-8.2* | Tau TuBulin Kinase 8.2 | WBGene00016541 |
| *ttx-1* | abnormal ThermoTaXis 1 | WBGene00006652 |
| *ttx-3* | abnormal ThermoTaXis 3 | WBGene00006654 |
| *twk-17* | TWiK family of potassium channels 17 | WBGene00006671 |
| *twk-2* | TWiK family of potassium channels 2 | WBGene00006657 |
| *twk-35* | TWiK family of potassium channels 35 | WBGene00006687 |
| *twk-39* | TWiK family of potassium channels 39 | WBGene00006690 |
| *twk-47* | TWiK family of potassium channels 47 | WBGene00018002 |
| *twk-48* | TWiK family of potassium channels 48 | WBGene00010784 |
| *twk-9* | TWiK family of potassium channels 9 | WBGene00006664 |
| *unc-10* | UNCoordinated 10 | WBGene00006750 |
| *unc-122* | UNCoordinated 122 | WBGene00006845 |
| *unc-17* | UNCoordinated 17 | WBGene00006756 |
| *unc-2* | UNCoordinated 2 | WBGene00006742 |
| *unc-29* | UNCoordinated 29 | WBGene00006765 |
| *unc-3* | UNCoordinated 3 | WBGene00006743 |
| *unc-30* | UNCoordinated 30 | WBGene00006766 |
| *unc-31* | UNCoordinated 31 | WBGene00006767 |
| *unc-36* | UNCoordinated 36 | WBGene00006772 |
| *unc-38* | UNCoordinated 38 | WBGene00006774 |
| *unc-41* | UNCoordinated 41 | WBGene00006777 |
| *unc-7* | UNCoordinated 7 | WBGene00006747 |
| *unc-75* | UNCoordinated 75 | WBGene00006807 |
| *unc-8* | UNCoordinated 8 | WBGene00006748 |
| *unc-86* | UNCoordinated 86 | WBGene00006818 |
| *unc-9* | UNCoordinated 9 | WBGene00006749 |
| *vab-3* | Variable ABnormal morphology 3 | WBGene00006870 |
| *W01B6.5* | Tyrosine-protein kinase | WBGene00012172 |
| *W02B12.12* | Protein kinase domain-containing protein | WBGene00012207 |
| *W03A5.1* | Protein kinase domain-containing protein | WBGene00020966 |
| *W03G9.5* | Protein kinase domain-containing protein | WBGene00021016 |
| *W06F12.3* | non-specific serine/threonine protein kinase | WBGene00012307 |
| *wht-4* | WHiTe (Drosophila) related ABC transporter 4 | WBGene00017179 |
| *wht-5* | WHiTe (Drosophila) related ABC transporter 5 | WBGene00008950 |
| *wht-8* | WHiTe (Drosophila) related ABC transporter 8 | WBGene00012925 |
| *xtr-2* | MX region of TRA-2 Related 2 | WBGene00006966 |
| *Y116A8C.24* | Tyrosine-protein kinase | WBGene00013801 |
| *Y32G9A.3* | Protein kinase domain-containing protein | WBGene00021302 |
| *Y38H8A.4* | Protein kinase domain-containing protein | WBGene00012638 |
| *Y39G8C.2* | Protein kinase domain-containing protein | WBGene00012731 |
| *Y43C5B.2* | Tyrosine-protein kinase | WBGene00012785 |
| *Y51H7C.9* | Serine racemase | WBGene00021787 |
| *Y52D5A.2* | Tyrosine-protein kinase | WBGene00021793 |
| *Y54F10BM.3* | Tyrosine-protein phosphatase domain-containing protein | WBGene00021858 |
| *Y65B4A.9* | Protein kinase domain-containing protein | WBGene00022032 |
| *Y69E1A.3* | Tyrosine-protein kinase | WBGene00013475 |
| *Y69F12A.1* | Protein kinase domain-containing protein | WBGene00022102 |
| *Y71F9AL.2* | Protein kinase domain-containing protein | WBGene00022108 |
| *Y71G12B.22* | Protein kinase domain-containing protein | WBGene00022157 |
| *Y75B8A.6* | CXXC-type domain-containing protein | WBGene00013543 |
| *ZC581.2* | non-specific serine/threonine protein kinase | WBGene00022632 |
| *ZC581.7* | Tyrosine-protein kinase | WBGene00022634 |
| *ZK177.2* | No official full name assigned | WBGene00022670 |
| *ZK354.2* | Protein kinase domain-containing protein | WBGene00022705 |
| *ZK354.6* | non-specific serine/threonine protein kinase | WBGene00022707 |
| *ZK507.1* | Putative serine/threonine-protein kinase ZK507.1 | WBGene00013978 |
| *ZK596.2* | non-specific serine/threonine protein kinase | WBGene00014007 |
| *ZK616.65* | Protein-tyrosine phosphatase | WBGene00235383 |
| *ZK622.1* | Tyrosine-protein kinase | WBGene00022780 |
| *zmp-1* | Zinc MetalloProtease 1 | WBGene00006987 |
| *ztf-30* | Zinc finger putative Transcription Factor family 30 | WBGene00015523 |
