## Supplementary Figures and Legends for "Cisplatin-triggered sensory dysfunction in NER-defective *C. elegans* involves GABAergic and cholinergic signaling"

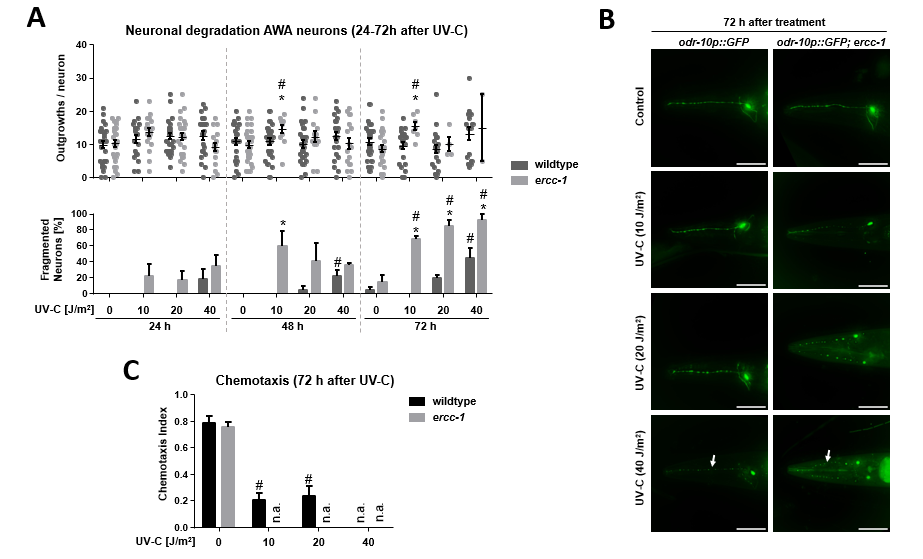

**Supplementary Fig. S1:** **UV-C induces AWA neuronal degeneration and sensory dysfunction in *ercc-1* mutants**

**A, B:** Quantification of neuronal degradation and damage (beading) of chemosensory AWA neurons in wildtype and *ercc-1* mutated nematodes 24 h, 48 h and 72 h after UV-C irradiation (0-40 J/m^2^) with representative images **(B)**. Scale bar = 50µm. Arrows indicate interruptions of the neuronal process interpreted as fragmentation/ neuronal cell death. Quantitative data shown are from n = 3 independent experiments, with at least 8 images being analyzed per experimental condition. *, p ≤ 0.05 (wildtype vs. mutant, students t-test); ^#^, p ≤ 0.05 (control vs. UV-C irradiated, One-Way ANOVA).

**C:** Chemotaxis analysis of *ercc-1* mutated animals 72 h after UV-C irradiation (0-40 J/m^2^). Data shown are the mean + SEM of three independent experiments (N ≥ 100 nematodes per experimental condition and experiment). ^#^, p≤ 0.05 (control vs. UV-C irradiated; One-Way ANOVA).

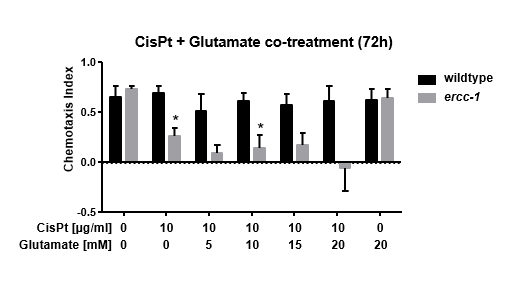

**Supplementary Fig. S2: Glutamate substitution does not alleviate CisPt-induced sensory dysfunction.**

Chemotaxis analysis of wildtype and *ercc-1* mutated animals after 72 h of co-treatment with CisPt (10 µg/ml) and glutamate (20 mM). Data shown are the mean + SEM of three independent experiments (N ≥ 100 nematodes per experimental condition and experiment). *, p ≤ 0.05 (wildtype vs. mutant, students t-test); ^#^, p ≤ 0.05 (control vs. CisPt-treatment, One-Way ANOVA).

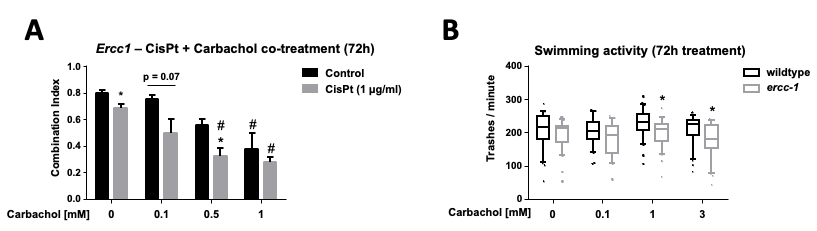

**Supplementary Fig. S3: Carbachol induces sensory dysfunction but not motoric dysfunction in *ercc-1* mutated animals**

**A:** Chemotaxis analysis of *ercc-1* mutated animals after 72 h of co-treatment with CisPt (1 µg/ml) and carbachol (0-1 mM). Data shown are the mean + SEM of three independent experiments (N ≥ 100 nematodes per experimental condition and experiment). *, p ≤ 0.05 (‑CisPt vs. +CisPt, students t-test); ^#^, p ≤ 0.05 (control vs. carbachol-treatment, One-Way ANOVA).

**B:** Swimming activity of wildtype and *ercc-1* mutated animals was analyzed after 72 h of carbachol treatment (0-3 mM). Data shown as boxplots with median (center line) box (25^th^-75^th^ percentiles) and whiskers (10^th^-90^th^ percentiles) of three independent experiments (N = 10 nematodes per experimental condition and experiment). *, p ≤ 0.05 (wildtype vs. mutant, students t-test); ^#^, p ≤ 0.05 (control vs. carbachol-treatment, One-Way ANOVA).

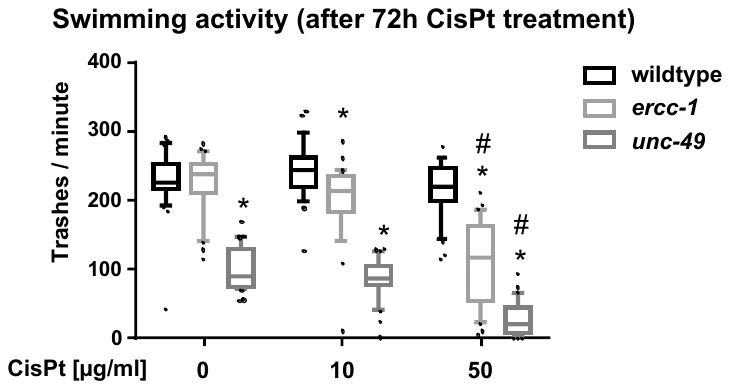

**Supplementary Fig. S4: Effects of CisPt on swimming activity of *ercc-1* and *unc-49* mutants.**

Swimming activity of wildtype, *ercc-1* and *unc-49* mutated animals after 72 h of CisPt treatment (0-50 µg/ml). Data shown as boxplots with median (center line) box (25^th^-75^th^ percentiles) and whiskers (10^th^-90^th^ percentiles) of three independent experiments (N = 10 nematodes per experimental condition and experiment). *, p ≤ 0.05 (wildtype vs. mutant, students t-test); ^#^, p ≤ 0.05 (control vs. carbachol-treatment, One-Way ANOVA).

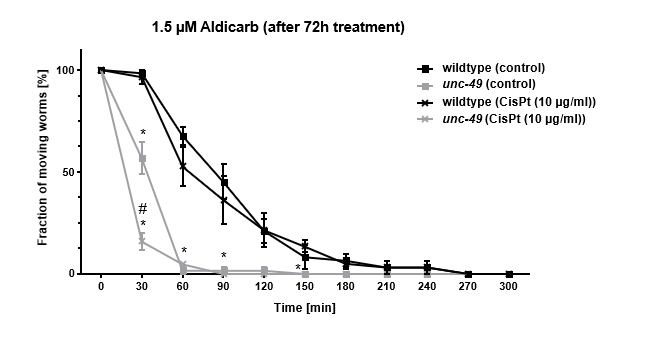

**Supplementary Fig. S5: GABAergic mutant exhibits increased sensitivity to aldicarb-induced paralysis, particularly following CisPt treatment.**

Aldicarb assay of wildtype and *unc-49* mutant animals after 72 h of CisPt treatment (0, 10 µg/ml) (N ≥ 60, n = 3). Data shown are mean ± SEM of three independent experiments with 18-21 nematodes being per experimental condition and experiment). *, p ≤ 0.05 (wildtype vs. mutant, students t-test); ^#^, p ≤ 0.05 (control vs. CisPt-treatment, One-Way ANOVA).

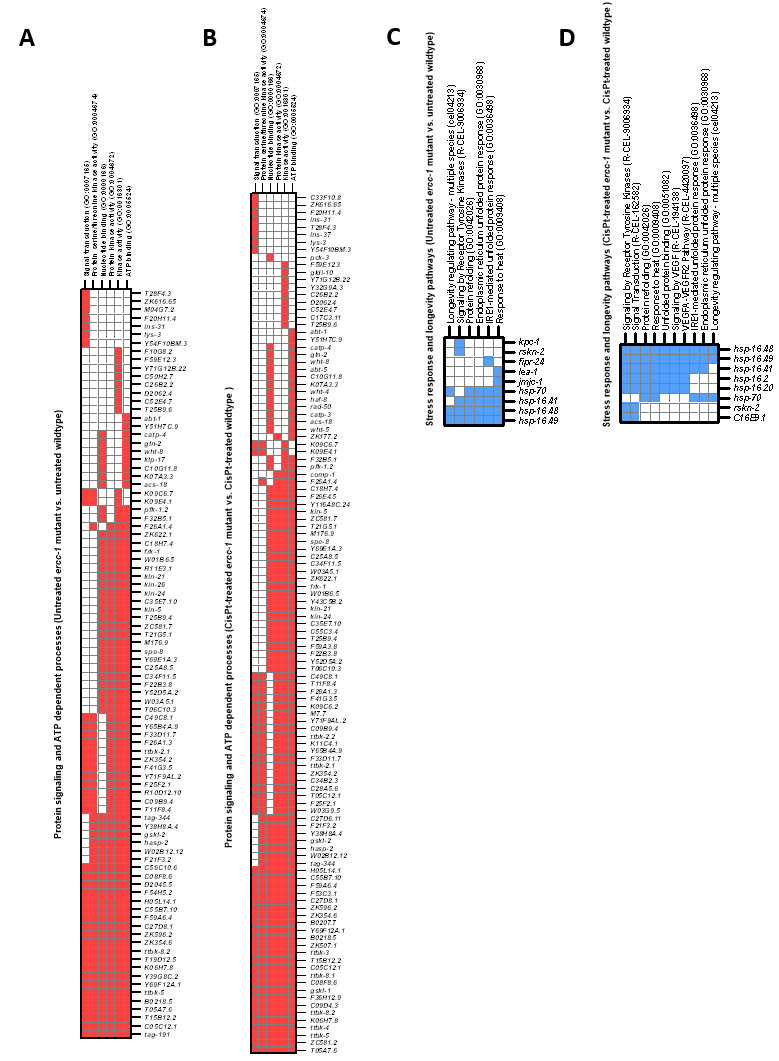

**Supplementary Fig. S6: Overview of shared functional annotation clusters in *ercc-1* mutants vs. wildtype animals under basal conditions and following CisPt-treatment.**

Composition of the functional annotation clusters presented in **Fig. 5**, showing the individual enriched terms and their associated differentially expressed genes. Terms were derived from GO, KEGG, and Reactome pathway analyses. Official WormBase names and identifiers are shown in **Supplementary Tab. S2**.

**A:** Protein signaling and ATP dependent processes (untreated *ercc-1* vs. untreated wildtype).

**B:** Protein signaling and ATP dependent processes (CisPt-treated *ercc-1* vs. CisPt-treated wildtype).

**C:** Stress response and longevity pathways (untreated *ercc-1* vs. untreated wildtype).

**D:** Stress response and longevity pathways (CisPt-treated *ercc-1* vs. CisPt-treated wildtype).

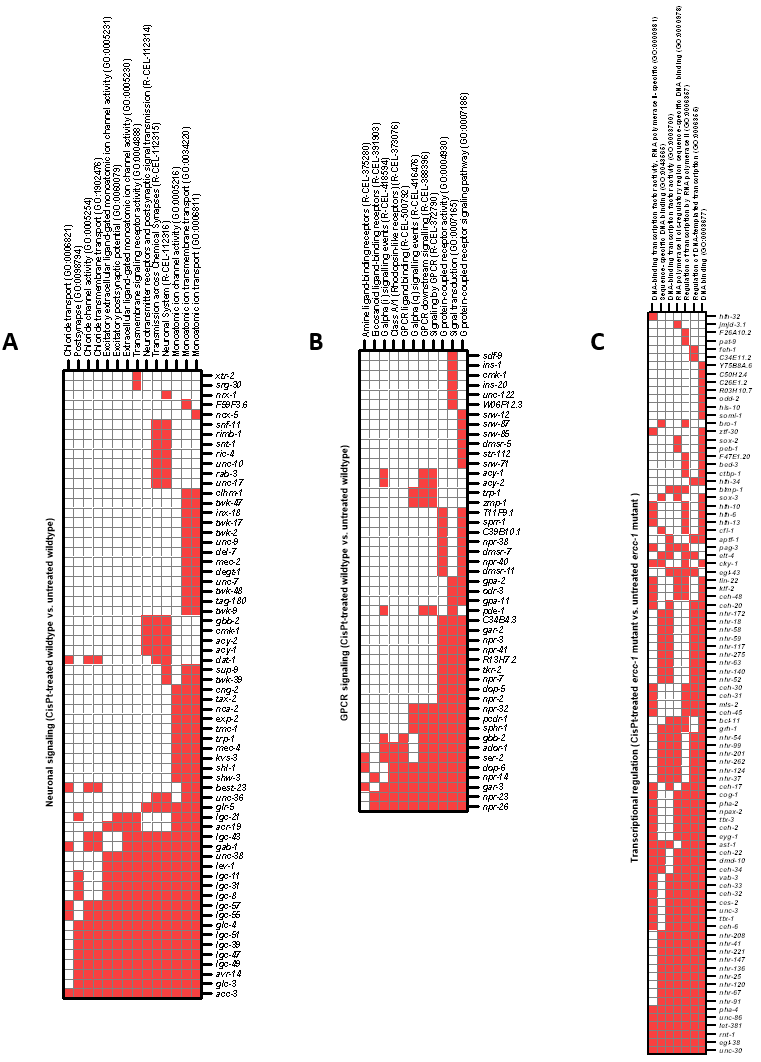

**Supplementary Fig. S7: Detailed overview of functional annotation clusters under basal conditions vs. CisPt-treatment in *ercc-1* mutants and wildtype animals.**

Composition of the functional annotation clusters presented in **Fig. 6**, showing the individual enriched terms and their associated differentially expressed genes. Terms were derived from GO, KEGG, and Reactome pathway analyses. Official WormBase names and identifiers are shown in **Supplementary Tab. S2**.

**A:** Neuronal signaling (CisPt-treated wildtype vs. untreated wildtype).

**B:** GPCR signaling (CisPt-treated wildtype vs. untreated wildtype).

**C:** Stress response and longevity pathways (CisPt-treated *ercc-1* vs. untreated *ercc-1*).

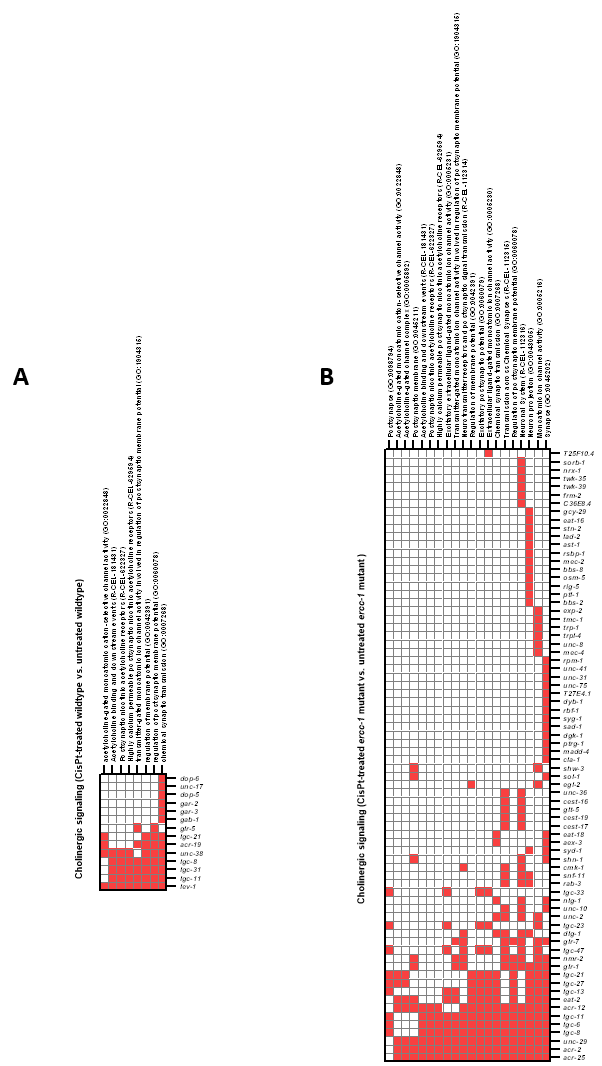

**Supplementary Fig. S8: Detailed overview of shared functional annotation cluster under basal conditions vs. CisPt-treatment in *ercc-1* mutants and wildtype animals.**

Composition of the functional annotation clusters presented in **Fig. 6**, showing the individual enriched terms and their associated differentially expressed genes. Terms were derived from GO, KEGG, and Reactome pathway analyses. Official WormBase names and identifiers are shown in **Supplementary Tab. S2**.

**A:** Cholinergic signaling (CisPt-treated wildtype vs. untreated wildtype).

**B:** Cholinergic signaling (CisPt-treated *ercc-1* vs. untreated *ercc-1*).
